# An intercellular metabolic relay for brain sparing in *Drosophila*

**DOI:** 10.1101/2025.04.02.645514

**Authors:** Adrien Franchet, Yuhong Jin, Clare L. Newell, Victor Girard, Gérard Manière, Yaël Grosjean, Christopher Barrington, James I. MacRae, Ian S. Gilmore, Alex P. Gould

## Abstract

Brain sparing protects growth of the central nervous system (CNS) at the expense of other developing organs during nutrient restriction. This survival strategy is conserved from *Drosophila* to mammals but little is known about its underlying metabolic mechanisms. Here, we show that CNS uptake and catabolism of circulating glutamine is essential for neural stem cell proliferation during brain sparing but not normal brain development. Glutamine is imported into perineurial glia of the blood-brain-barrier, which upregulate glutaminase (Gls) and vesicular glutamate transporter 1 (VGlut1) to promote glutaminolysis and glutamate secretion. Neural stem cells then import released glutamate via excitatory amino acid transporter 1 (Eaat1) and maintain it at homeostatic levels via glutamine synthetase 1 (Gs1). Glutamine nitrogen contributes surprisingly little to newly synthesized DNA or protein in neural stem cells but its carbon fuels the oxidative tricarboxylic acid cycle. These findings identify an intercellular metabolic relay specifically required to sustain the proliferation of neural stem cells during brain sparing.

## Introduction

Developing animals of most, if not all, species adjust their growth trajectories in response to decreases in nutrient availability. Nutrient deprivation, unless severe or early onset, is usually compatible with development into undersized yet viable and fecund adults. Interestingly, however, not all organs scale down isometrically with the body. In particular, allometric growth of the central nervous system (CNS) is prioritized over that of other developing organs, a process known as brain sparing^1–3^. Brain sparing exists in humans, rodents, insects and likely other animals and is thought to be a key strategy for surviving growth restriction^1–7^. In mammals, brain sparing during fetal development is known to involve the redistribution of blood flow from abdominal visceral organs towards the developing CNS^8,9^. Nevertheless, molecular mechanisms that function in neural stem cells and/or in their local microenvironment, the niche^10–15^, to support brain sparing have yet to be identified in mammals.

The CNS of the insect *Drosophila* is a well characterised model system with powerful genetic tools^16–21^. In the adult *Drosophila* brain it is known that energy homeostasis and cell survival are promoted via glial-neuronal lactate and alanine shuttles similar to those of mammals^22–27^. Less is known about how metabolic shuttles operate during development of the *Drosophila* CNS, when neural stem cells, called neuroblasts, divide asymmetrically to generate diverse neuronal and glial subtypes^28–34^. During the larval phase of development, each neuroblast and its progeny become ensheathed within a chamber formed by cortex glia^35–37^. The cortex glia, together with astrocyte-like glia, ensheathing glia and also the perineurial and subperineurial glia constitute a neural stem cell niche providing neuroblasts with cell contacts and signals that regulate their proliferation^5,38–46^. The perineurial and subperineurial glia form the blood-brain-barrier (BBB), which regulates the exchange of metabolites and ions between the circulation (hemolymph) and the CNS^47–50^. Fuelled by glycolysis and oxidative phosphorylation, neuroblast divisions accelerate during larval development and can attain a speed of about one hour per cell cycle ^51–53^.

The *Drosophila* larval CNS is a tractable system for identifying cellular and molecular mechanisms involved in brain sparing^5–7,54^. In the contexts of hypoxia or oxidative stress that is strong enough to inhibit epithelial progenitor proliferation almost completely, neuroblast cell divisions are spared by 30% to 100%, depending upon the conditions^7^. A key protective metabolic response of the larval CNS to hypoxia and oxidative stress is the induction of lipid droplets in cortex and subperineurial glia^7,55^. Glial synthesis of lipid droplet triacylglycerols can promote neuroblast sparing during oxidative stress, although the non-cell autonomous mechanism involved is not known^7^. In the context of strong nutrient restriction (NR), which prevents all larval weight gain, neuroblasts are nevertheless able to sustain their cell divisions at very close to the normal speed^4–6^. This impressive brain sparing requires anaplastic lymphoma kinase (Alk) signalling within neuroblast lineages^5^. Unlike nutrient-dependent insulin-like receptor (InR) signalling^41,42,56^, Alk signaling is able to sustain neuroblast proliferation during NR at mid-to-late larval stages^5^. Alk also promotes sparing of the major endocrine gland in *Drosophila* and its ectopic expression in an organ that is not normally spared during NR is sufficient to activate sparing^5,57^. Emerging evidence indicates that there are metabolic changes in glia of the neural stem cell niche during NR. For example, the carbohydrate transporter Tret1-1 is upregulated in perineurial glia^58^ and the orphan solute carrier Pathetic is required in glia for the normal diameter of the brain lobes^59^. It is, however, not clear how specific amino acids are metabolised during NR via the various glial subtypes of the BBB and neural stem cell niche, nor how they may impact neural stem cell proliferation during brain sparing. Here, we use nutrient supplementations to identify specific circulating amino acids that support brain sparing. Focusing on one non-essential amino acid fuel that we find is necessary for brain sparing, its metabolism is tracked using stable isotope tracing, mass spectrometry imaging and a genetically-encoded metabolite sensor. Cell-type specific genetic manipulations of neural stem cells, neurons and glial subtypes are then used to identify the first, to our knowledge, intercellular metabolic pathway dedicated to neural stem cell sparing during nutrient restriction.

## Results

### Glutamine supports neuroblast proliferation *ex vivo*

We first measured amino acid (AA) levels in a *Drosophila melanogaster* model of brain sparing, in which neural stem cell (neuroblast) proliferation is maintained close to 100%^5^. Larvae of a control strain, *w^1118^ iso31* (Ref 60), were raised on a standard yeast-based diet (STD) until they reached ∼1 mg, which occurs at ∼66 hr after larval hatching (ALH), and then switched to nutrient restriction (NR) for a further 24 hr (**Fig. 1a**). The application of NR at this stage ensures that critical weight (CW) has been attained and therefore that development is not delayed by this dietary manipulation^5,61–63^. AAs were quantified in the developing CNS and in the circulation (cell-free hemolymph) at 66 hr (CW) and also at 90 hr ALH in fed and NR larvae (**Fig. 1b,c**). During development from 66 to 90 hr on STD, there is a general tendency for the levels of essential and non-essential AAs to increase in both the hemolymph and the CNS. Following 24 hr of NR, however, many AAs show moderate to strong decreases in the in the CNS and also in the hemolymph. For the CNS, there were particularly strong reductions of 53% for alanine (Ala), 77% for proline (Pro), 90% for threonine (Thr), and 48% for glutamate (Glu), compared to age-matched STD larva (**Fig. 1b,c**, and **Table 1**). Importantly, for the metabolically related amino acids glutamine (Gln) and Glu, high concentrations of the former but not the latter are present in the hemolymph, whereas this relationship is reversed in the CNS. Glu is a major central and peripheral neurotransmitter in insects^64,65^ and, as in mammals, it is kept at a low concentration in the circulation^66,67^. The AA analysis shows that Ala, Pro, Thr and Gln/Glu become substantially depleted in the CNS, and hemolymph, during *in vivo* NR and are therefore candidates for metabolic fuels involved in the growth of spared organs including the CNS.

**Figure 1:**
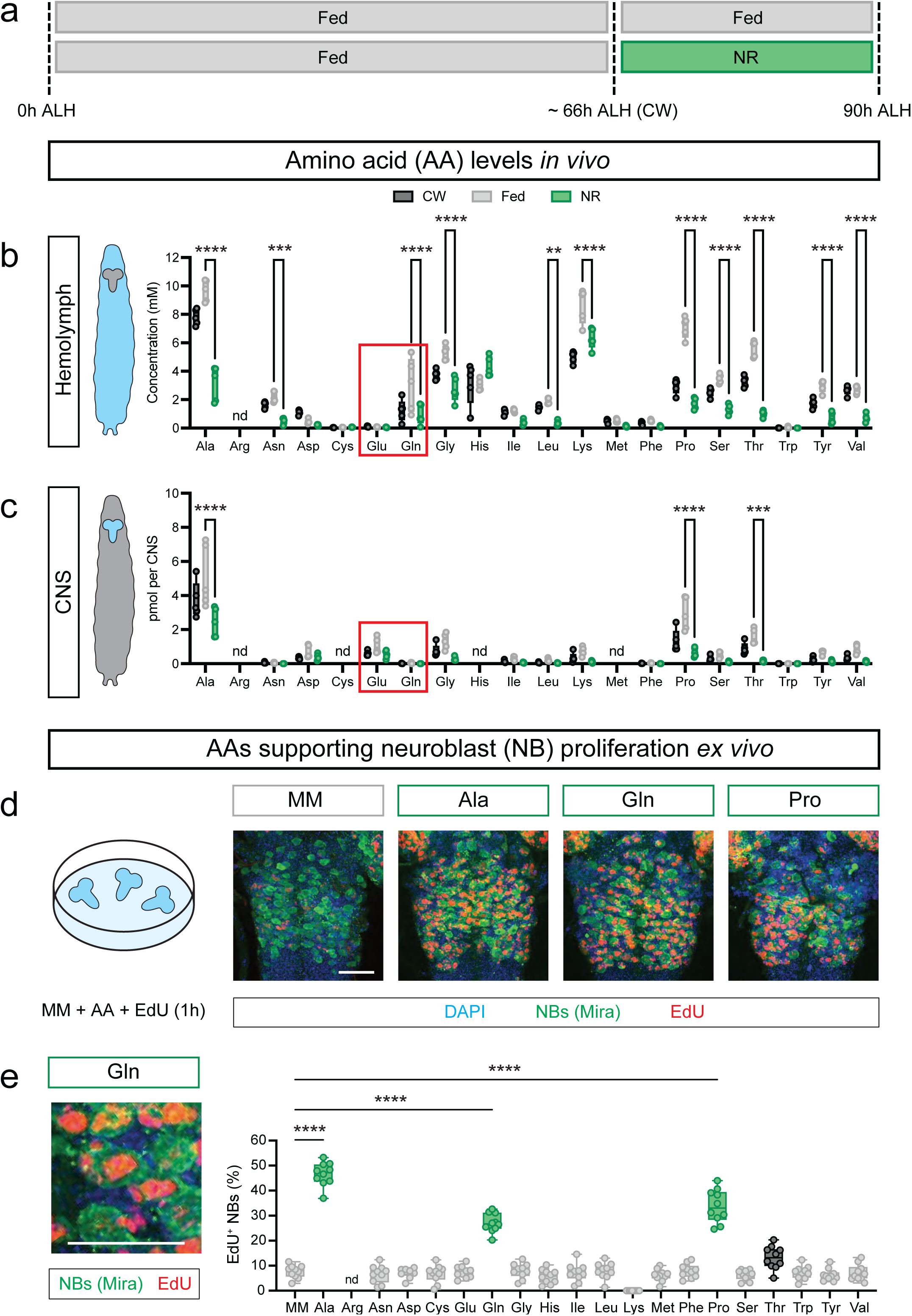
Glutamine, proline and alanine support *ex vivo* brain sparing. **a**, The *Drosophila* model for brain sparing. Timeline shows larval development from 0 to 90 hours after larval hatching (ALH), highlighting critical weight (CW) at ∼66h and showing the Fed conditions for normal development and the nutrient restriction (NR) paradigm for brain sparing. **b**-**c**, *In vivo* amino acid levels in larval hemolymph (mM) (b) and larval CNS (pmol) (c) are shown for CW, Fed and NR larvae. **d-e**, *Ex vivo* short-term cell proliferation assays quantifying EdU incorporation into neuroblasts (Mira^+^ cells) in the absence (MM) or presence of the indicated amino acid concentration as in **Table 1**. Representative confocal images are shown at low (d) or high (e) magnifications, along with quantifications for each of the 20 amino acids. Ala, alanine; Arg, arginine; Asn, asparagine; Asp, aspartate; Cys, cysteine; Glu, glutamate; Gln, glutamine; Gly, glycine; His, histidine; Ile, isoleucine; Leu, leucine; Lys, lysine; Met, methionine; Phe, phenylalanine; Pro, proline; Ser, serine; Thr, threonine; Trp, tryptophan; Tyr, tyrosine; Val, valine. Scale bar = 50 µm. Statistical analysis: One-way ANOVA with multiple comparisons; \**p* < 0.05 (black), \*\*\*\**p* < 0.0001 (green).

To test the potential contributions of Ala, Pro, Thr, Gln/Glu and other amino acids to brain sparing, we used a short-term assay for neural stem cell proliferation. An *ex vivo* set up was chosen as this allows the direct effects of each amino acid upon the CNS to be assessed without complications from metabolic processes in other tissues. Whole larval CNSs were incubated in minimal medium (MM) alone, or with the addition of one amino acid, and the percentage of type I thoracic neuroblasts incorporating 5-ethynyl-2’-deoxyuridine (EdU) into DNA was quantified^7,29,30^. With a 3 hr incubation in MM alone, only ∼5-10% of neuroblasts passed through S-phase during the last hour (**Fig. 1d,e**). This was also the case for 17 of the 20 proteogenic amino acids added at their physiological concentrations in fed hemolymph (**Fig. 1d,e**). In contrast, physiological concentrations of Ala, Pro or Gln were sufficient to sustain EdU incorporation in ∼30-50% of neuroblasts during the 1 hr labelling period (**Fig. 1d,e**). Equalizing each amino acid at an artificial concentration of 10 mM gave similar results for Ala, Pro and Gln but now Glu, and to a lesser extent Thr, were also able to maintain short-term EdU incorporation (**Extended Data Fig. 1**). The analysis thus far provides evidence that Ala, Pro, and Gln/Glu are abundant in hemolymph, depleted from the CNS during *in vivo* brain sparing and that each is sufficient to sustain *ex vivo* neuroblast proliferation in the acute brain sparing assay.

### Glutamine atoms contribute little to neuroblast synthesis of DNA, proteins or fatty acids

Three reasons make it interesting to focus on Gln/Glu metabolism and how this sustains brain sparing. First, glutamine is an abundant non-essential AA with many potential roles in carbon and nitrogen metabolism^68–74^. Second, it is intriguing that glutamine is at a high concentration in the hemolymph but barely detectable within the CNS, yet the converse is the case for glutamate. And third, two key enzymes of glutamine metabolism, glutaminase and glutamine synthase, each catalyse opposite and predominantly unidirectional reactions *in vivo*^71,75^. This allows loss-of-function phenotypes to be attributed to one direction of the enzyme reaction, which is not possible with bidirectional enzymes.

Isotope tracing was utilized to identify which of the many potential pathways of glutamine metabolism^76–78^ are relevant within the developing CNS. We substituted unlabelled for isotope labelled Gln using the *ex vivo* conditions where Gln is sufficient to sustain neuroblast proliferation in MM over 3 hr. The duration of the incubation is too short to reach isotopic steady-state in the medium but longer *ex vivo* incubations are prone to CNS deterioration. To determine the Gln nitrogen contribution to macromolecule biosynthesis, OrbiSIMS mass spectrometry imaging was used at ambient temperature on fixed tissue sections^79^. This detects strong signals from adenine fragmented and ionised from nucleic acid but not the free nucleobase, and from Gln or Glu fragmented and ionised from proteins but not free amino acids^79–82^. It is challenging to measure absolute metabolite concentrations using mass spectrometry imaging^83–85^ but isotopic fractional enrichments can be accurately determined. Using Gln-amide-^15^N_1_ or Gln-amine-^15^N_1_, the frequency distributions of fractional enrichments in 5 μm pixels were determined globally across the CNS and also specifically in CNS progenitors. To achieve this cell-type specific metabolic imaging, the medium was supplemented with a low concentration (10 µM) of ^13^C_10_^15^N_5_-deoxyadenosine, which marks neural progenitors (neuroblasts and ganglion mother cells, GMCs) via salvage pathway incorporation into newly-synthesised DNA, detected as the ^13^C_5_^15^N_5_-adenine ion (*m/z=*144.049). However, labelling with Gln-amide-^15^N_1_ or Gln-amine-^15^N_1_ gave little or no detectable enrichment of adenine M+1 (*m/z=*135.044) in neural-progenitor specific or global populations of pixels (**Extended Data Fig. 2a,b**). Isotopic enrichments of Gln M+1 and Glu M+1 isotopologues in protein were also very low in the vast majority of neural progenitor or global pixels. Similarly, global measurements via gas chromatography mass spectrometry (GC-MS) detected little or no incorporation of carbon from ^13^C_5_-Gln into the major fatty acids of the larval CNS (palmitate C16:0, palmitoleate C16:1, stearate C18:0 and oleate C18:1) (**Extended Data Fig. 2c**). Together, these results demonstrate that glutamine contributes very little carbon or nitrogen to newly synthesized DNA, proteins or fatty acids during *ex vivo* brain sparing.

### CNS glutamine fuels anaplerosis and biosynthesis of amino acids and neurotransmitters

We next used GC-MS to quantify the incorporation of nitrogen from labelled Gln into small polar metabolites (**Fig. 2a**). CNSs incubated with ^15^N_2_-Gln showed high fractional enrichments for the M+1 isotopologues of Glu (0.97), GABA (0.94), aspartate (0.85) and alanine (0.79) (**Fig. 2b** and **Table 2**). This shows that the CNS imports Gln and metabolises it via amidohydrolysis and aminotransferase reactions. CNSs were also incubated with ^13^C_5_-Gln to characterise the relative contributions of glutamine carbon to the oxidative and reductive branches of the tricarboxylic acid (TCA) cycle^86–88^ (**Fig. 2c**). The M+4 isotopologue of Glu (the ion measured contains only four of the five Glu carbons) had a fractional enrichment of 0.58 and M+4 GABA had a fractional enrichment of 0.92, indicative of Gln amidohydrolysis and Glu decarboxylase reactions, respectively (**Fig. 2d** and **Table 2**). For TCA cycle intermediates, the fractional enrichments of the M+4 isotopologues (Suc 0.59, Fum 0.50) were much higher than those of their corresponding M+3 isotopologues (Suc 0, Fum 0.02) (**Fig. 2d**). This shows that the CNS preferentially metabolizes Gln carbon via the oxidative rather than the reductive TCA cycle. In summary, the surprising conclusion of the combined isotope imaging and tracing experiments is that Gln can sustain neuroblast cell cycles while contributing very little carbon or nitrogen to the synthesis of DNA, fatty acids or proteins. Instead, the CNS imports Gln to fuel the oxidative TCA cycle, as well as neurotransmitter biosynthesis and aminotransferase reactions.

**Figure 2:**
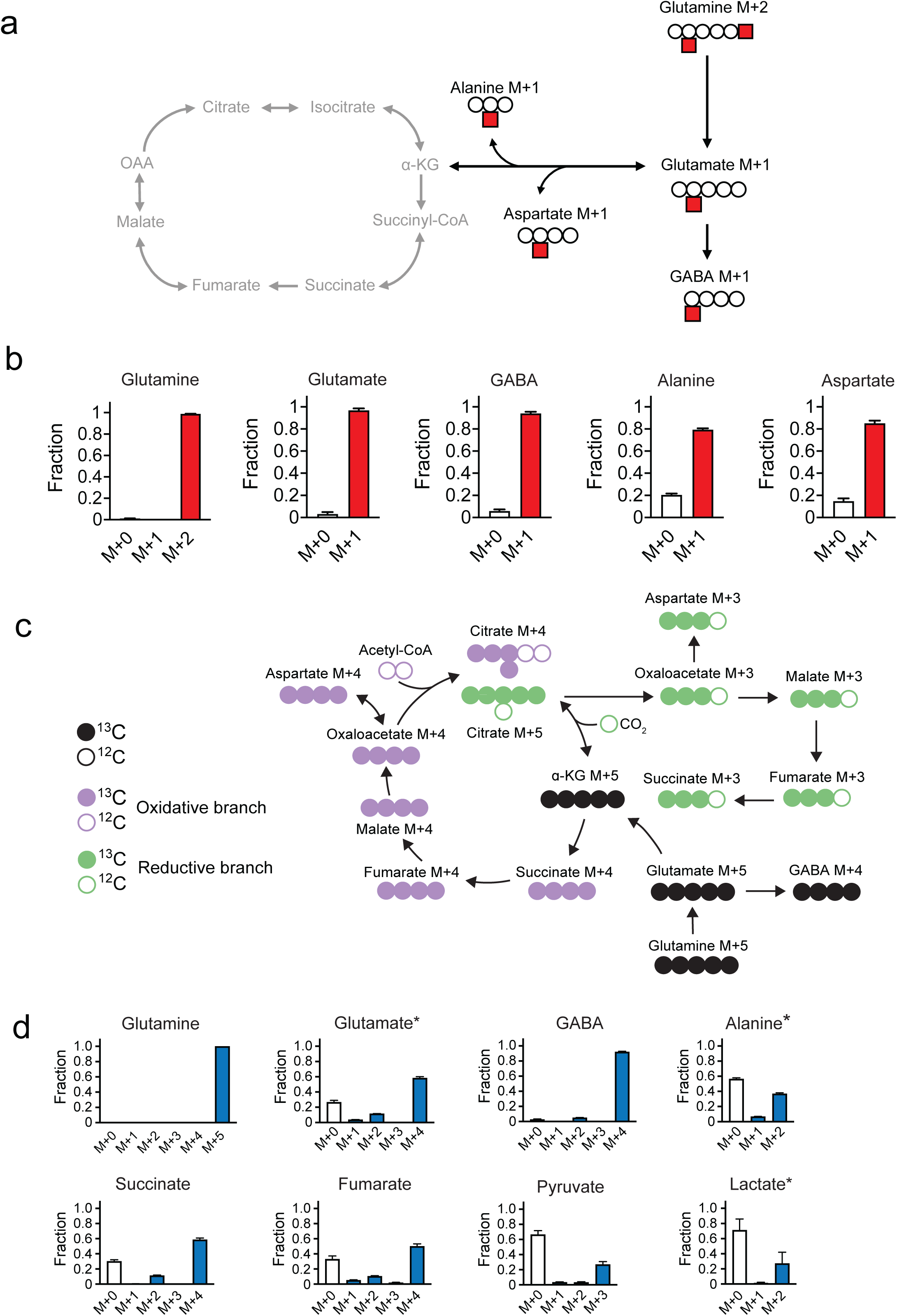
Glutamine contribute to the oxidative branch of the TCA cycle. **a**, Potential metabolic fates of ^15^N from ^15^N_2_-glutamine. **b**, GC-MS fractional enrichments of isotopologues of glutamine, glutamate, GABA, alanine, and aspartate in larval CNSs incubated for 3 hr in minimal medium supplemented with ^15^N_2_-glutamine. **c**, Isotopologues obtained when ¹³C₅-glutamine (glutamine M+5) is metabolized via the oxidative or reductive branches of the TCA cycle. **d**, Fractional enrichments for metabolites obtained from isotope tracing of *ex vivo* CNSs incubated in minimal medium supplemented with ¹³C₅-glutamine for 3 hr. * indicates that, for glutamate, alanine and lactate the measured ions contain one less carbon than the intact molecule.

### Brain sparing requires glutaminolysis in perineurial glia and glutamine biosynthesis in neuroblast lineages

To identify *in vivo* roles for Gln/Glu metabolism during brain sparing, we analyzed a conserved glutamine amidohydrolase (glutaminase, Gls) and a conserved glutamate-ammonia ligase (glutamine synthetase, Gs1) (**Fig. 3a**). To guide subsequent genetic analysis of specific cell types within the neuroblast lineage and its glial niche (**Fig. 3b**), we first conducted single-nucleus RNA sequencing of 27,794 cells in the CNS at 90 hr ALH (**Extended Data Fig. 3a**). Using previously validated cell-type specific markers, strong expression of Gs1 was observed in neurons and Gls in perineurial glia of the blood-brain barrier (BBB), with weaker Gls expression in immature and mature neurons (**Extended data Fig. 3b)**. We also used a protein trap allele (*Gls^MI09647-GFSTF.0^*) to show that Gls is expressed strongly in the BBB as well as in mature larval neurons and, importantly, it is also upregulated during NR-induced brain sparing (**Extended data Fig. 4a,b**).

**Figure 3:**
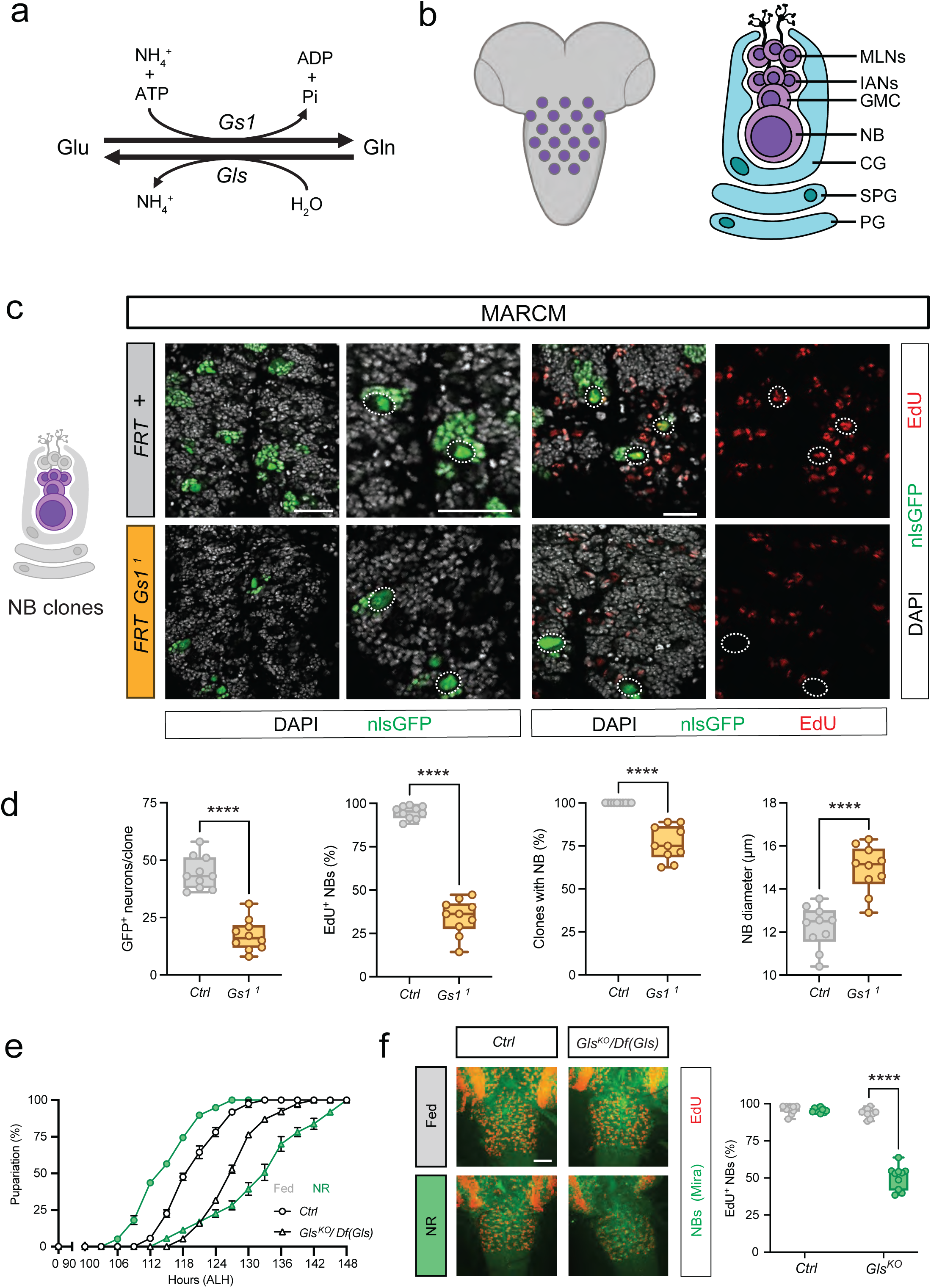
*Gs1* and *Gls* are required for neuroblast proliferation. **a**, The glutamine synthetase 1 (Gs1) and glutaminase (Gls) enzyme reactions. **b**, Schematic of neuroblasts within the larval CNS (left) and the cellular components (right) of the neuroblast lineage (purple) and its niche glia (blue). Perineural glia (PG) and subperineurial glia (SPG) form the blood-brain-barrier, cortex glia (CG) surrounds the neuroblast lineage, which is composed of neuroblasts (NBs), ganglion mother cells (GMC), immature adult neurons (IANs) and mature larval neurons (MLNs). **c**, Neuroblasts incorporate less EdU in *Gs1^1^* mutant FRT than in control FRT MARCM clones (marked with nlsGFP). **d**, *Gs1^1^* mutant FRT clones have deficits in the number of GFP+ neurons per clone and in the percentage of neuroblasts incorporating EdU. They also lack a neuroblast in ∼25% of clones and it is enlarged in the remaining 75% clones. **e**, Time-to-pupariation curves show that Gls^KO^ mutant (Gls^KO^*/Df(2R)Excel7121*) larvae are moderately delayed compared to controls (*y w*) and they lack the NR-induced developmental acceleration of controls. **f**, EdU incorporation in neuroblasts of Gls^KO^*/Df(2R)Excel7121* larvae, compared to controls (*y w*), is similar in fed larvae but strongly decreased in NR larvae. Scale bar = 50 µm. Statistics used unpaired t-tests and two-way ANOVA with multiple comparisons. \*\*\*\**p* < 0.0001.

The functions of *Gs1* and *Gls* during brain sparing were determined *in vivo* using mutant alleles and RNAi knockdowns. In particular, we sought to characterise the fed versus NR requirements for *Gs1* and *Gls* in each of the cell types of the neuroblast lineage and its glial niche (**Fig. 3b**). To characterise the loss-of-function phenotype of *Gs1* in type I neuroblast lineages of the thoracic region, we used mosaic analysis with a repressible cell marker (MARCM)^89^ to make neuroblast lineage clones homozygous for *Gs1^1^*, a lethal amorphic allele^90,91^. *Gs1^1^* clones generate an average of only ∼17 GFP^+^ daughter cells whereas control clones contain ∼44 GFP^+^ cells (**Fig. 3c,d** and **Extended Data Videos 1 and 2**). This cell number deficit correlates with a pronounced decrease in neuroblast proliferation, loss of a recognizably large neuroblast in ∼24% of clones and an abnormally enlarged neuroblast in the remaining ∼76% of clones (**Fig. 3c,d**). Hence, during fed development, *Gs1* is required within neuroblast lineages for cell proliferation and the maintenance of normal stem cell size. For *Gls,* we characterised a knockout allele, *Gls^CR70270-KO-kG4^* (hereafter called *Gls^KO^*), where coding sequence has been replaced with a TI{KozakGAL4} cassette^92^. *Gls^KO^* is larval viable when hemizygous over a deficiency, with only a minor delay to pupariation of ∼6 hr (**Fig. 3e**). In addition, the modest NR-induced acceleration to pupariation, characteristic of *Drosophila* (Ref 61), is lost in *Gls^KO^*mutants and, instead, there is a delay of ∼18 hr compared to controls (**Fig. 3e**). Importantly, EdU incorporation by type I neuroblasts is decreased by ∼50% in *Gls^KO^*mutants but only in the context of NR (**Fig. 3f**). Thus, *Gls* activity is required for neuroblast proliferation but only during brain sparing.

We next used the GAL4/UAS system^93^ to pinpoint which cell types in the larval CNS require *Gs1* and *Gls in vivo*. Consistent with the phenotype of *Gs1^1^* clones, RNAi knockdown of *Gs1* specifically in neuroblast lineages (*nab-GAL4*) strongly decreased EdU incorporation in both fed and NR neural stem cells (**Fig. 4a**). In contrast to neuroblast lineage knockdown, pan-glial (*repo-GAL4*) inactivation of *Gs1* did not decrease neuroblast EdU incorporation during either fed or NR (**Fig. 4b**). Neuroblast lineage knockdown of *Gs1* decreased the size of some neuroblasts, as we had observed in ∼25% of *Gs1^1^* MARCM clones, and the DNA in these stem cells was abnormally condensed (**Extended Data Fig. 5a**). In addition, *ex vivo* supplementations of the medium with Gln, Glu, or Thr were unable to rescue neuroblast proliferation in *Gs1* knockdowns (**Fig. 4c**). Further investigation revealed that some *Gs1*-deficient neuroblasts activate the effector caspase Dcp-1 *in vivo* and are thus likely to be undergoing apoptosis (**Extended Data Fig. 5b**). To pinpoint the cell type(s) within the neuroblast lineage that require *Gs1*, we knocked it down either in mature (*nSyb-GAL4*) or immature (*nerfin-1-GAL4*) neurons, or in both, but EdU incorporation was not affected (**Fig. 4d**). Currently available genetic tools are not able to restrict loss-of-function specifically to only the neuroblast and/or the GMC. Nevertheless, together with the strong MARCM and *nab-GAL4* RNAi phenotypes, these findings suggest that *Gs1* is required in neural progenitors (neuroblasts and/or GMCs) for the appropriate size, proliferation and survival of the stem cell. One caveat here is that we cannot distinguish whether *Gs1* is solely required in neural progenitors, where snRNAseq suggests that the expression is very low, or whether there might be redundant requirements in progenitors and in immature neurons. Either way, it is clear that *Gs1* is required for functional neuroblasts and that its role is constitutive rather than specific to brain sparing.

**Figure 4:**
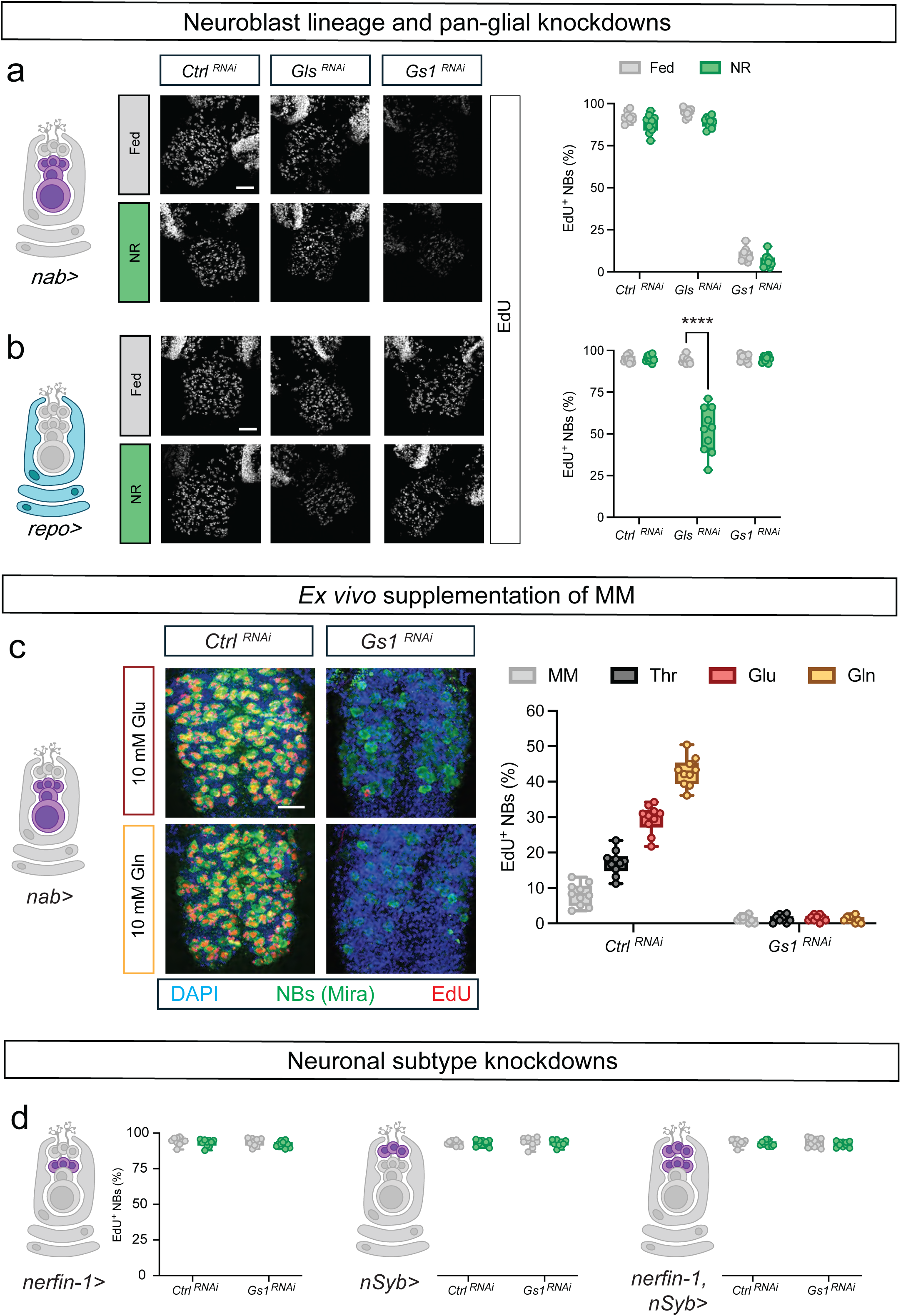
Constitutive neuroblast *Gs1* and NR-specific glial *Gls* functions. **a**, *In vivo* neuroblast EdU incorporation following knockdowns in neuroblasts and immature adult neuronal progeny is significantly altered during fed and NR conditions for *Gs1* (*nab>Gs1^RNAi^*) but not for *Gls* (*nab>Gls^RNAi^*) or for controls (*nab>mCherry^RNAi^*). **b**, *In vivo* neuroblast EdU incorporation following pan-glial knockdowns is significantly altered during NR but not fed conditions for *Gls* (*repo>Gls^RNAi^*) but not for *Gs1* (*repo>Gs1^RNAi^*) or for controls (*repo>mCherry^RNAi^*). **c**, Neuroblasts of *ex vivo* CNSs incubated in minimal medium (MM) supplemented with Thr, Glu or Gln incorporate EdU in controls (*nab>mCherry^RNAi^*) but not *Gs1* knockdowns in neuroblasts and immature adult neuronal progeny (*nab>Gs1^RNAi^*). **d**, *In vivo* neuroblast EdU incorporation following *Gs1* knockdowns in immature adult neurons (*nerfin-1>Gs1^RNAi^*), mature larval neurons (*nSyb>Gs1^RNAi^*) or both (*nerfin-1,nSyb>Gs1^RNAi^*) are not significantly altered during fed or NR conditions. Controls are *nerfin-1>mCherry^RNAi^*, *nSyb>mCherry^RNAi^*, and *nerfin-1,nSyb>mCherry^RNAi^*. Scale bar = 50 µm. Statistical tests were two-way ANOVA with multiple comparisons; \*\*\*\**p* < 0.0001.

For *Gls*, *in vivo* knockdown in neuroblast lineages did not significantly decrease fed or NR proliferation of neural stem cells, which is in marked contrast to *Gs1* (**Fig. 4a**). Importantly, however, pan-glial knockdown of *Gls* specifically decreased NR but not fed neuroblast incorporation by ∼50% (**Fig. 4b**). Along with the *Gls^KO^* results, these findings reveal an *in vivo* glial requirement for *Gls* that is specific for brain sparing. Pan-glial *Gls* knockdown also decreases substantially the ability of 1 mM or 4 mM Gln to stimulate *ex vivo* neuroblast proliferation (**Fig. 5a**). This key finding demonstrates that glial *Gls* activity is required to mediate the pro-proliferative effect of Gln on neuroblasts. Nevertheless, a supraphysiological concentration of 10 mM Gln is able to bypass the glial *Gls* deficit (**Fig. 5a**). Similarly, *in vivo* supplementation of the NR medium with 5 mM glutamate, 10 mM glutamine or both rescues, to varying extents, the loss of brain sparing observed with pan-glial *Gls* knockdown, but this effect could be mediated via metabolism in multiple organs not just the CNS (**Fig. 5b**). Given that high but not low concentrations of Gln can bypass the *Gls* proliferation phenotype, we reasoned that there could be partial redundancy with another glial enzyme that metabolizes glutamine into glutamate. Consistent with this, the phenotype of a widely expressed glutamine-hydrolyzing NAD synthase (*Nadsyn*) mimicks that of *Gls*, such that pan-glial but not neuroblast-lineage knockdown suppresses NR proliferation (**Extended data Fig. 3b** and **Extended data Fig. 8a,b**). To determine which glial subtype is responsible for the neuroblast sparing phenotype of *Gls*, it was knocked down specifically in cortex (*Cyp4g15-GAL4*), subperineurial (*moody-GAL4*) or perineurial (*NP6295-GAL4*) glia. This revealed that knockdown in perineurial glia recapitulates the NR-specific phenotype of a ∼50% decrease in neuroblast proliferation *in vivo* (**Fig. 5c**). Importantly, knockdown in perineurial glia, as in all glia together, significantly blunted the ability of 1 mM Gln to stimulate *ex vivo* neuroblast proliferation (**Fig. 5d**). Hence, *Gls* is required in perineurial glia for neuroblast proliferation when the CNS is exposed to physiological concentrations of glutamine. With our previous finding that perineurial glia upregulate *Gls* during NR, the genetic analysis and AA supplementations demonstrate clearly that glutaminolysis in the perineurial glia of the BBB is required specifically for brain sparing.

**Figure 5:**
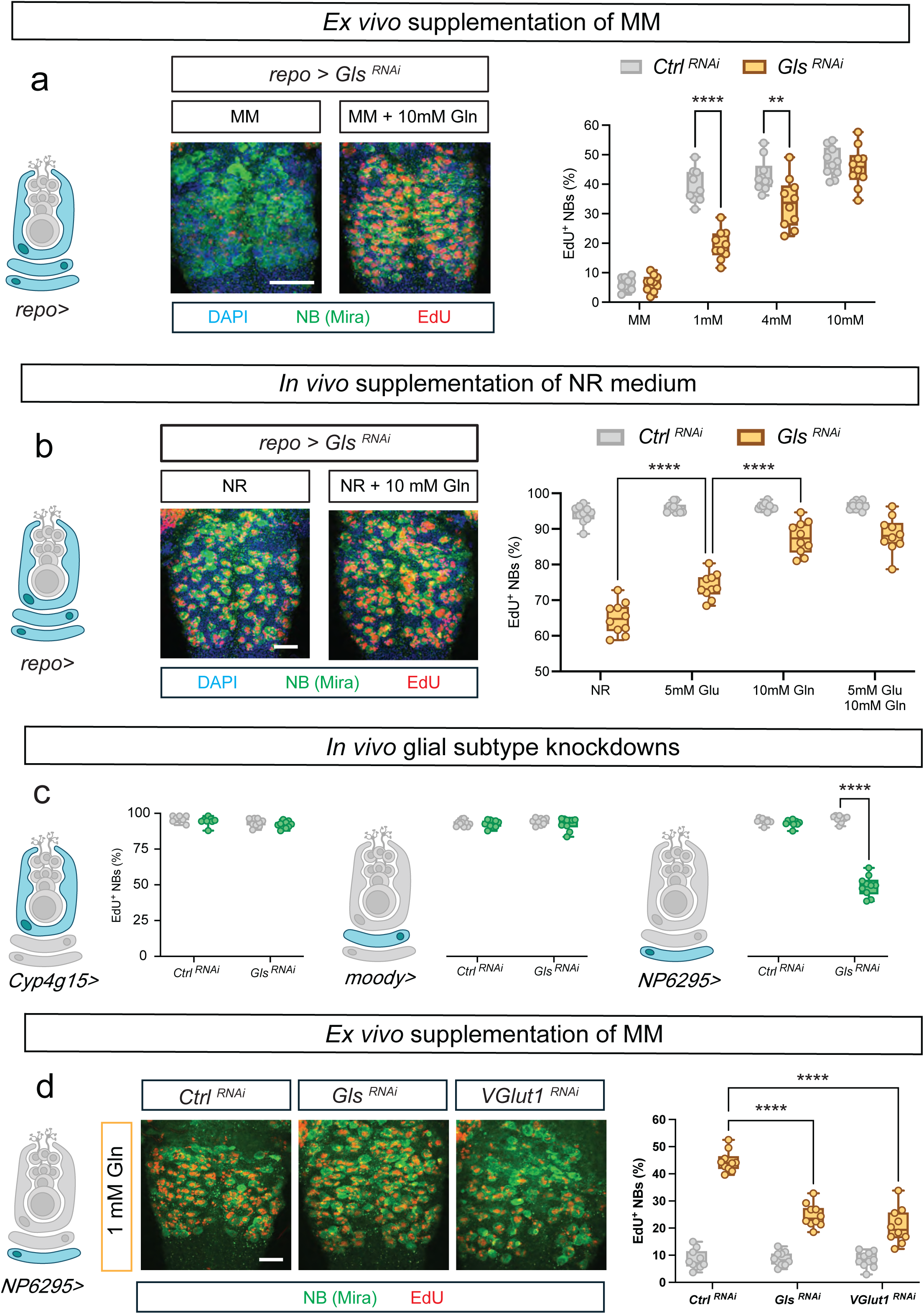
*Gls* in perineurial glia mediates glutamine-dependent neuroblast proliferation. **a**, Glutamine stimulation of *ex vivo* neuroblast EdU incorporation is compromised following pan-glial Gls knockdown (*repo>Gls^RNAi^*) at 1 mM and 4 mM but not at 10 mM. **b**, *In vivo* neuroblast EdU incorporation during NR is compromised by pan-glial *Gls* knockdown (*repo>Gls^RNAi^*) but partially rescued via dietary supplementation with 5 mM Glu, 10 mM Gln or both. **c**, *In vivo* neuroblast EdU incorporation during NR is substantially decreased by *Gls* knockdown in perineurial glia (*NP6293>Gls^RNA^*) but not cortex glia (*Cyp4g15>Gls^RNA^*) or subperineurial glia (*moody>Gls^RNA^*). **d**, 1 mM glutamine stimulation of *ex vivo* neuroblast EdU incorporation is significantly compromised following knockdown in perineural glia of *Gls* (*NP6293>Gls^RNA^*) or *VGlut1* (*NP6293>VGlut1^RNA^*). Scale bar = 50 µm. Statistical tests used two-way ANOVA with multiple comparisons; \*\*\*\**p* < 0.0001, \*\**p* < 0.01.

### Glial *Gls* and neuroblast lineage *Gs1* ensure glutamate homeostasis of stem cells

To monitor local extracellular glutamate, we utilized a circularly permutated GFP sensor that is membrane bound, iGluSnFR.A184A (Ref 94). First, we characterised this glutamate sensor using live imaging in *ex vivo* CNS cultures. Static imaging of BBB glia expressing iGluSnFR indicates a dose-dependent response to glutamate added to the medium (**Fig. 6a**). Time-lapse recordings revealed that iGluSnFR expressed in neuroblast lineages responded to exogenous glutamine within 60 seconds (**Fig. 6b)**. These observations, together with our previous isotope tracing experiments, demonstrate that glutamine enters BBB glia and is converted into glutamate, which accumulates in the extracellular space inside the cortex glial chambers where the neuroblasts reside. Importantly, *Gs1* knockdown in postembryonic neuroblast lineages leads to a strong and sustained iGluSnFR signal increase in these cells, detectable using either static or time-lapse modes of live imaging (**Fig. 6c,d)**. This shows that *Gs1* is required in neuroblast lineages to prevent the accumulation of excessive glutamate in the extracellular space inside cortex glial chambers.

**Figure 6:**
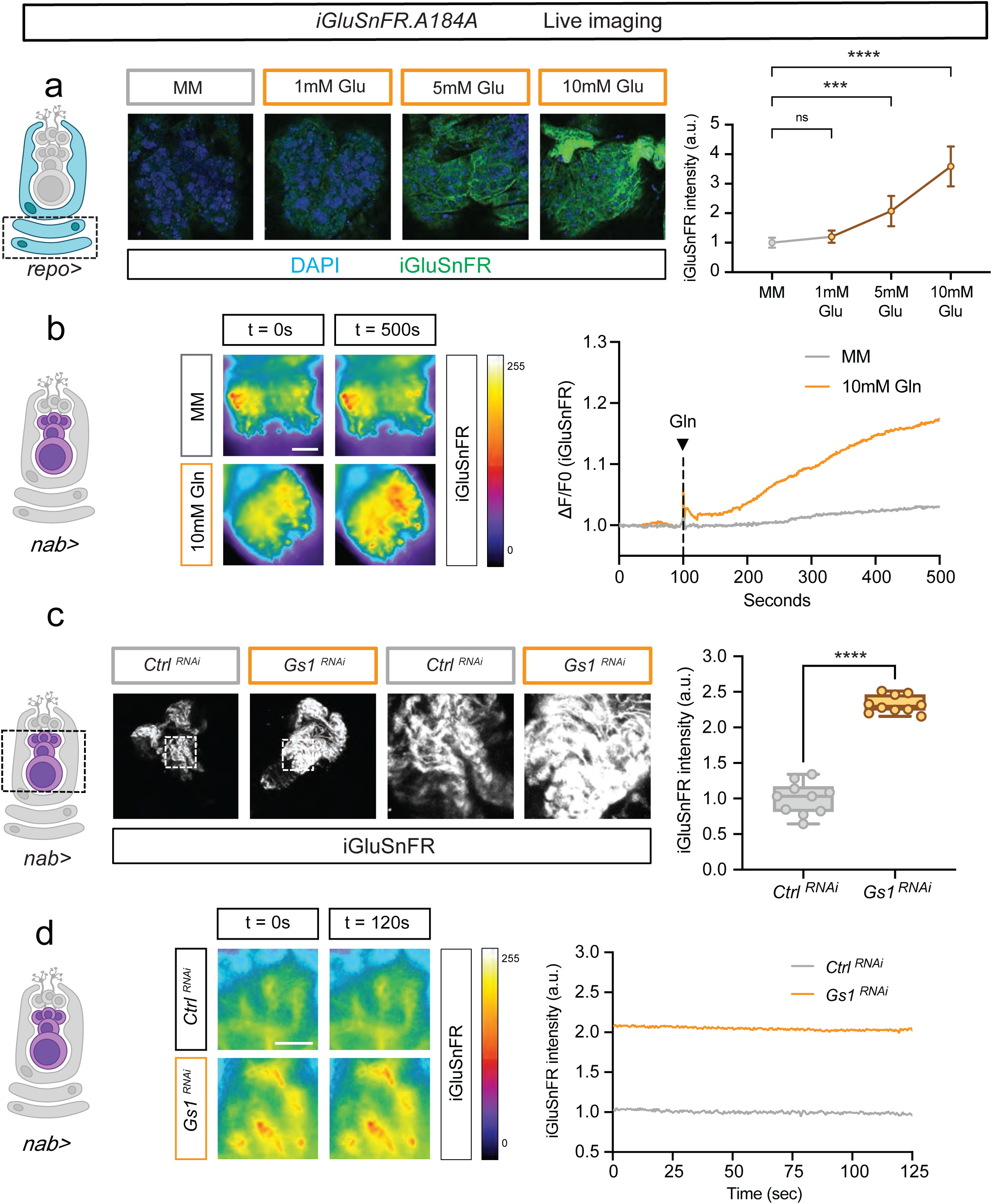
Neuroblast lineage *Gs1* prevents extracellular glutamate accumulating in cortex glial chambers. **a**, Live static imaging of *ex vivo* CNSs expressing the glutamate sensor iGluSnFR in glia (*repo>iGluSnFR*), incubated in minimal medium (MM) supplemented with 1 mM, 5 mM, or 10 mM Gln. The dose-response curve (right) shows iGluSnFR signal intensity versus Gln concentration. **b**, Live time-lapse imaging of *ex vivo* CNSs expressing iGluSnFR in neuroblasts and immature adult neuronal progeny (*nab>iGluSnFR*), incubated in minimal medium (MM) with 10 mM Gln at 100 seconds (s). Panels (left) show the iGluSnFR signal at 0 s and 500 s and the graph (right) shows the iGluSnFR signal change (ΔF/F0) versus time. **c**, Live static imaging of *ex vivo* CNSs co-expressing iGluSnFR in neuroblasts and immature adult neuronal progeny with a control RNAi (*nab>iGluSnFR, Ctrl^RNAi^*) or with *Gs1* RNAi (*nab>iGluSnFR, Gs1^RNAi^*). iGluSnFR signals in both genotypes are shown in CNSs at low and high magnification (left) and quantified graph (right). **d**, Live time-lapse imaging of *ex vivo* CNSs co-expressing iGluSnFR in neuroblasts and immature adult neuronal progeny with a control RNAi (*nab>iGluSnFR, Ctrl^RNAi^*) or with *Gs1* RNAi (*nab>iGluSnFR, Gs1^RNAi^*). iGluSnFR signals in both genotypes are shown in CNSs at 0 s and 120 s (left) and quantified in the graph (right) over a 125 s period. Scale bar = 50 µm. Statistical analysis: Two-way ANOVA with multiple comparisons; \*\*\*\**p* < 0.0001, \*\*\**p* < 0.001.

It is technically challenging with live imaging to monitor iGluSnFR with the spatial resolution achievable using confocal analysis of fixed material. We noticed, however, that the extracellular glutamate increase following *Gs1* knockdown in neuroblast lineages could be detected with iGluSnFR.A184A not only in live imaging mode but, surprisingly, also after mild fixation (**Extended data Fig. 6a**). This corresponds to a higher ratio of iGluSnFR to anti-GFP fluorescence, strongly suggesting that it reflects *bona fide* glutamate signal rather than sensor protein expression (**Extended data Fig. 6a**). It is not yet clear how iGluSnFR signal is retained after fixation but we note that the A184A version has a higher affinity and slower off rate for glutamate than the A184V variant used to detect millisecond changes in neuronal glutamate^94–96^. Importantly, the iGluSnFR.A184A signal at the plasma membranes of neuroblasts increased strongly in fixed CNSs from *Gs1* knockdown larvae raised in fed and NR conditions (**Extended data Fig. 6b**). In contrast, following glial *Gls* knockdown, iGluSnFR.A184A signal at the plasma membranes of cortical glia, including those juxtaposed to neuroblasts, significantly decreased but only during NR (**Extended data Fig. 6c**). Hence, glial *Gls* is required specifically during NR for glutamate to accumulate to physiological levels in the extracellular spaces inside cortex glial chambers, where the neuroblasts reside.

### Glutamate is exported from PG via VGlut1 and imported into NB lineages via Eaat1

We next addressed the mechanism by which glutamate is transported during brain sparing from the perineurial glia of the BBB to the cortex glial chambers. Two evolutionarily conserved transporters of glutamate in *Drosophila* are excitatory amino acid transporter 1 (Eaat1) and vesicular glutamate transporter 1 (VGlut1). *Drosophila* Eaat1 is a sodium-dependent importer of glutamate known to be expressed and required in glia^97–99^. *Drosophila* VGlut1 is an exporter of glutamate from neurons, loading it into synaptic vesicles for release during neurotransmission^65,100,101^.

*Eaat1* is expressed in cortex and astrocyte-like glia, and also in neuroblasts but not in most neurons (**Extended data Fig. 3b** and **Extended data Fig 7a**). Expression of *Eaat1* has also been detected in FACS purified neuroblasts^102^. Pan-glial *Eaat1* knockdown using a validated RNAi transgene is lethal, indicating an essential role but precluding larval analysis (**Fig. 7a**). In contrast, neuroblast lineage knockdown of *Eaat1* is larval viable but has a strong underproliferation phenotype that is specific for NR (**Fig. 7b**). Importantly, neuroblast lineage *Eaat1* is required for glutamate-dependent proliferation of neuroblasts *ex vivo* (**Fig. 7c**). Nevertheless, as with glial *Gls* knockdown, 10 mM glutamine was able to bypass this requirement (**Fig. 7c**).

**Figure 7:**
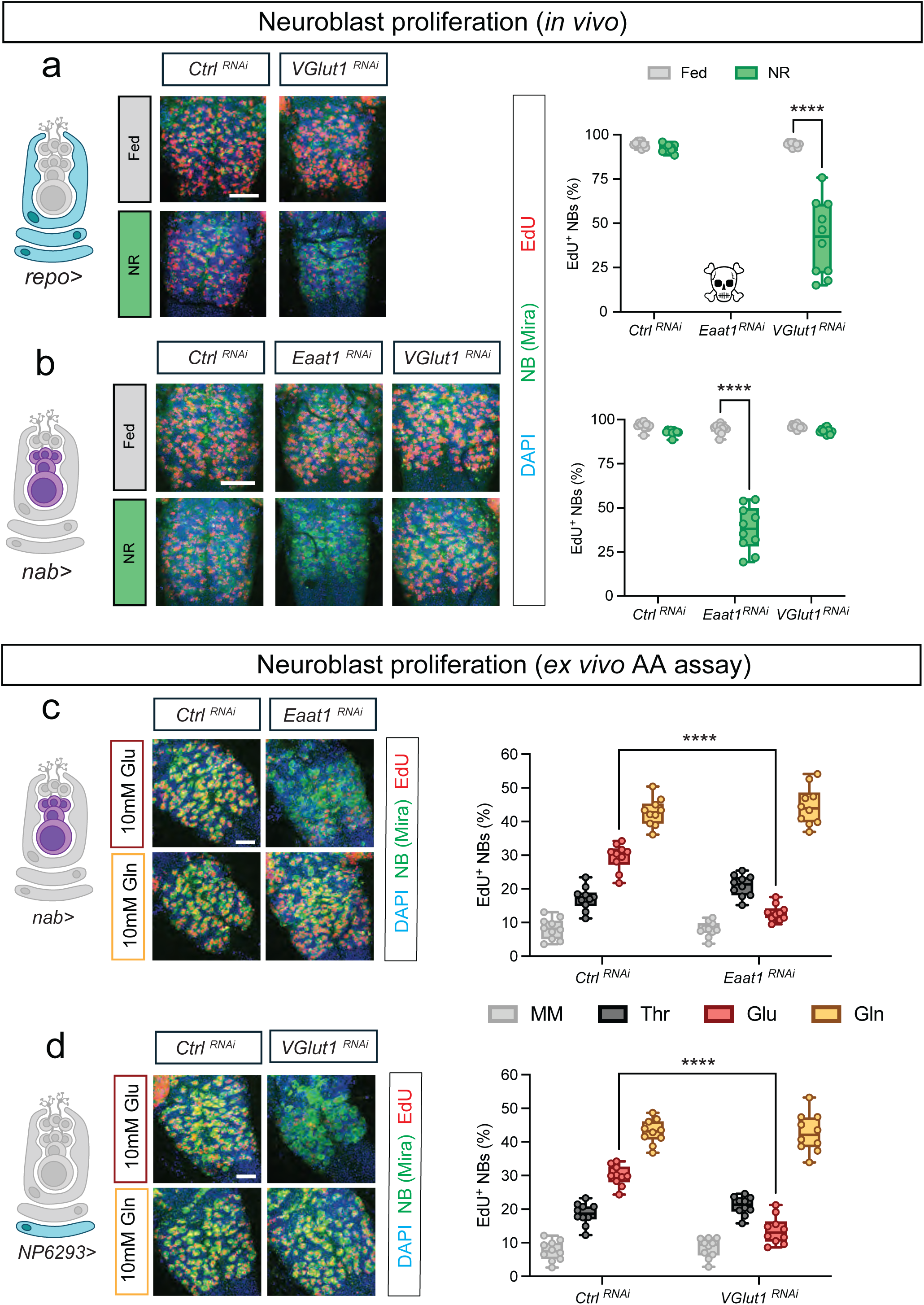
VGlut1 in perineurial glia and Eaat1 in neuroblast lineages mediate glutamate-dependent neuroblast proliferation. **a**, *In vivo* neuroblast EdU incorporation following pan-glial knockdown of *VGlut1* (*repo>VGlut1^RNAi^*) is substantially decreased during NR but not fed conditions. Pan-glial knockdown of *Eaat1* (*repo>Eaat1^RNAi^*) was lethal. **b**, *In vivo* neuroblast EdU incorporation following *Eaat* knockdown in neuroblasts and immature adult neuronal progeny (*nab>Eaat1^RNAi^*) is substantially decreased during NR but not fed conditions. Similar knockdown of *VGlut1* (*nab>VGlut1^RNAi^*) did not significantly alter neuroblast EdU incorporation compared with genetic controls (*nab>Ctrl^RNAi^*). **c**-**d**, Neuroblasts of *ex vivo* CNSs incubated in minimal medium (MM) supplemented with Thr, Glu or Gln. The ability of 10 mM Glu but not Thr or Gln to stimulate neuroblast EdU incorporation is significantly decreased following *Eaat* knockdown in neuroblasts and immature adult neuronal progeny (*nab>Eaat1^RNAi^*) (c), or following *VGlut1* knockdown in perineurial glia (*NP6293>VGlut1^RNAi^*) (d). Scale bar = 50 µm. Statistical analysis: Two-way ANOVA with multiple comparisons; \*\*\*\**p* < 0.0001.

*VGlut1* is highly expressed in the mature larval and immature adult neurons of the fed larval CNS (**Extended data Fig. 3b**). *VGlut1^OK371^*–*GAL4* recapitulates *VGlut1* expression^103^ and drives reporter expression in glutamatergic neurons of fed larvae (**Extended data Fig. 7b**). Interestingly, *VGlut1^OK371^*–*GAL4* expression is upregulated during NR, not only in neurons but also in perineurial glia of the BBB (**Extended data Fig. 7b,c**). Pan-glial but not neuroblast lineage knockdown of *VGlut1 in vivo*, using a validated RNAi transgene, lead to a variable but NR-specific deficit in neuroblast proliferation (**Fig. 7a,b**). Furthermore, *ex vivo* supplementations showed that perineurial glial activity of *VGlut1* is required for 1 mM glutamine, and even for 10 mM glutamate, to stimulate neuroblast proliferation (**Fig. 5d** and **Fig. 7d**). The *in vivo* and *ex vivo* results together demonstrate that both neuroblast *Eaat1* and perineurial *VGlut1* are required specifically for brain sparing during NR. They also strongly suggest that, during brain sparing, glutamate is exported from perineurial glia via VGlut1 and imported from the extracellular space of the cortex glial chamber into neuroblasts via Eaat1.

## Discussion

This study reveals the first mechanism in any species, to our knowledge, dedicated to brain sparing rather than to brain development in general. Central to the sparing mechanism is the role of glutamine/glutamate metabolism, which safeguards the proliferation of neural stem cells specifically during nutrient restriction. We now highlight how two glutamine/glutamate metabolic enzymes (Gs1 and Gls) and two glutamate transporters (Eaat1 and VGlut1) work together as a metabolic intercellular relay for brain sparing. We also discuss the functions of glutamine in neural progenitors and highlight striking contrasts between the cellular routes that metabolise it during brain sparing versus neurotransmission.

### A glutamine-glutamate intercellular relay dedicated to brain sparing

*In vivo* and *ex vivo* genetic and iGluSnFR analyses, together with stable isotope tracing data, support a metabolic relay model of brain sparing (**Fig. 8a**). Perineurial glia of the BBB take up circulating glutamine and convert it into glutamate via Gls. They export glutamate via the vesicular transporter VGlut1 and it accumulates within cortex glial chambers, where neuroblast lineages reside. Extracellular glutamate is then imported from the cortex glial chambers into neuroblasts via Eaat1. Within neuroblast lineages, glutamate feeds the oxidative TCA cycle and sustains neural stem cell proliferation specifically during brain sparing. Gs1 in neuroblast lineages ensures metabolic homeostasis, preventing excess accumulation of glutamate and perhaps ammonia. Although Gs1 plays a constitutive role in neuroblast proliferation, the other components of the intercellular relay (Gls, VGlut1 and Eaat1) are only required for this process during brain sparing. Moreover, at least two of these (VGlut1 and Gls) are also upregulated during brain sparing. Four other key findings underpin the glutamine-glutamate relay model for brain sparing. First, the developing CNS readily imports glutamine and efficiently metabolizes it into glutamate, which is used to feed the TCA cycle as well as to synthesize other amino acids. Second, the ability of glutamine/glutamate to stimulate neural stem cell proliferation requires Gls and VGlut1 in perineurial glia and also Gs1 and Eaat1 in neuroblast lineages. Third, Gs1 in neuroblast lineages prevents excess extracellular glutamate from accumulating inside cortex glial chambers. And fourth, glial activity of Gls is required, only during brain sparing, to sustain physiological levels of extracellular glutamate inside cortex glial chambers, where neuroblasts reside. Future studies will reveal how the glutamine-glutamate intercellular relay is integrated with the Alk signalling pathway, which functions as a constitutive pro-proliferative input for neural stem cells during brain development and sparing^5^.

**Figure 8:**
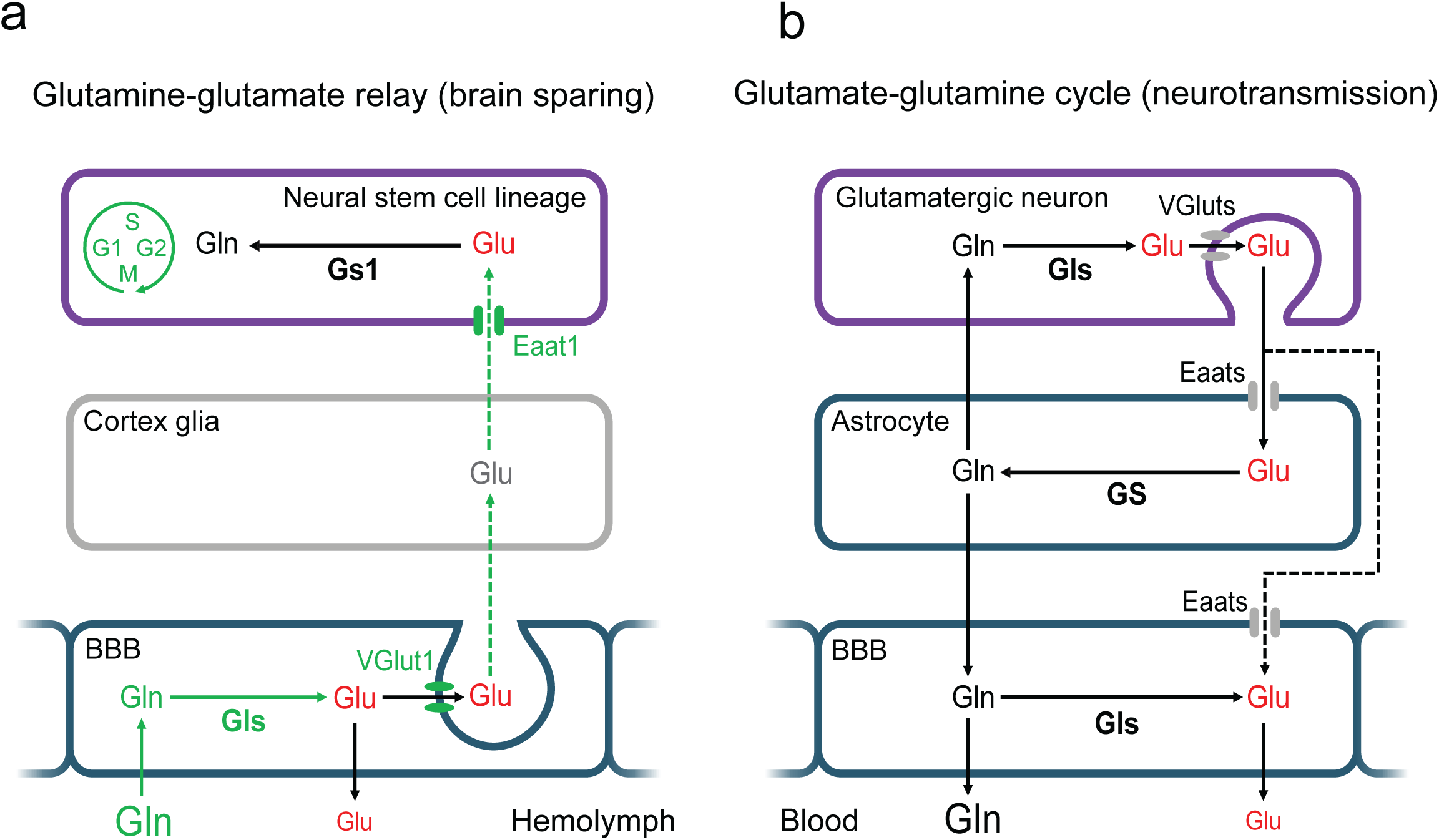
Glutamine metabolism during brain sparing versus neurotransmission. **a**, *Glutamine-glutamate intercellular relay model for brain sparing* In *Drosophila*, glutamine (Gln) in the circulation (hemolymph) is taken up by perineurial glia of the blood-brain barrier (BBB) and converted into glutamate (Glu). Glutamate is then released from perineurial glia and transported into the extracellular space inside the cortex glial chamber containing the neural stem cell and its daughter cells (neural stem cell lineage). Neural stem cells and perhaps their progeny import glutamate to feed the oxidative TCA cycle and to drive the cell cycle (G1, S, G2, M) during brain sparing. The conversion of some glutamate back into glutamine ensures metabolic homeostasis of the neural stem cell lineage. Genetic components dedicated to brain sparing but not required for normal CNS development are indicated in green. **b**, *Extended glutamate-glutamine cycle for neurotransmission* In mammals, glutamate neurotransmitter released from glutamatergic neurons is cleared from the synaptic cleft by astrocytes and converted into glutamine, which is recycled back into glutamate in neurons and exported from the CNS via endothelial cells of the BBB. Note that in both *Drosophila* and mammals, circulating glutamine is at a much higher concentration than glutamate. Glutamine is converted into glutamate via glutaminase (Gls) and glutamate is converted into glutamine via glutamine synthetase (GS, Gs1). Glutamate is exported from cells via vesicular glutamate transporters (VGluts, VGlut1) and imported into cells via excitatory amino acid transporters (Eaats, Eaat1).

The external location of perineurial glia of the BBB places them in an ideal position to sense circulating nutrients. Consistent with this, we showed that they respond to nutrient restriction by upregulating molecular machinery for glutamine utilization (Gls) and transport (VGlut1). This identifies an important function for perineurial glia as a gatekeeper of amino acid metabolism during brain sparing. As perineurial glia are also known to upregulate a trehalose transporter (Tret1) upon starvation^58^, a gatekeeping role in sparing neural stem cell proliferation might also extend to carbohydrate metabolism. Future studies will be required to decipher the role of other glial subtypes in brain sparing and whether or not they participate in the glutamine-glutamate relay.

### Glutamine fuels the oxidative TCA cycle during brain sparing

This study developed a quantitative method of mass spectrometry imaging to achieve spatial isotope tracing of metabolites at 5 μm resolution. We combined this technique with a mass tag for newly synthesized DNA in order to calculate isotope enrichments of metabolites specifically within neural progenitors. It is surprising that, over the time course of several glutamine-dependent neuroblast cycles, little nitrogen from this amino acid gets incorporated into DNA or protein in dividing progenitors and not much carbon is routed into fatty acids either. Instead, CNS glutamine predominantly fuels the oxidative TCA cycle as well as the biosynthesis of amino acids.

### Wiring brain glutamine metabolism to support neural stem cells versus neurons

In the mammalian brain, a glutamate-glutamine cycle prevents excitotoxicity via astrocytic recycling of glutamate neurotransmitter released from neurons ^74,104,105^. This neuron-astrocyte metabolic coupling works together with the BBB to export excess glutamate and glutamine from the brain into the blood^106–108^ In adult *Drosophila*, the available evidence thus far suggests that similar glutamate-glutamine metabolism may mediate glial support of neuronal activity^109–111^. It is striking that all four genetic elements (Gls, Gs1, Eaat1 and VGlut1) of the *Drosophila* glutamine-glutamate relay for brain sparing have mammalian orthologues that participate in the extended glutamate-glutamine cycle (**Fig. 8b**). However, these components and the cells that express them are wired together with very different metabolic logics. This reflects “opposite” functions of the two pathways - one supplies glutamate to neural stem cells and the other removes excess glutamate from neurons (**Fig. 8a,b**). A specific cellular difference is that neural stem cell lineages convert excess glutamate into glutamine during brain sparing whereas this role is carried out by astrocytes during neurotransmission. Another distinction is that VGluts export glutamate from the BBB during brain sparing but release it from neurons in the adult brain. In mammals, VGlut family transporters are active in neurons and have not been reported in the BBB, although recent evidence suggests they may function in a specialized subset of astrocytes^112^. In summary, glutamine metabolism is wired differently between cell types in the brain, depending upon whether it functions primarily to spare the proliferation of neural stem cells or to maintain the activity of neurons.

## Methods

### *Drosophila* husbandry and strains

Fly lines were raised on our standard diet (Fed) with a 12 hr light/dark cycle at 25°C with 60% humidity. Standard diet contains 58.5 g/L glucose, 66.3 g/L cornmeal, 23.4 g/L dried yeast, 7.02 g/L agar, 1.95 g/L Nipagen, and 7.8 mg/L Bavistan. Flies were left to lay eggs for 2 hours, on plates containing grape juice agar with yeast paste in the centre^60^. *Drosophila* strains used in this study were a Wolbachia-negative. In this study, we generated these *Drosophila* lines: *FRT40, Gs1^1^* mutant / nab-Gal4, UAS-iGluSnFR / repo-Gal4, UAS-iGluSnFR / nerfin-1-Gal4, nSyb-Gal4. We also tested two RNAi lines giving weak or no phenotypes: Gs1 RNAi (BL #52974) & Eaat1 RNAi (BL #43287). All *Drosophila* lines used for this study are listed in the **Table 3**.

### Larval Dietary Manipulations

For larval collection, flies raised at 25°C on our standard diet were transferred on a cage with grape juice agar plate plus yeast paste for 3 hours. After egg maturation for 24 hours, hatched L1 larvae were collected from the agar plates during a 1 hour time window using blunt forceps, and 50 individuals were transferred to each vial at 25°C on standard diet and raised to critical weight (∼1 mg at 60 hours after larval hatching) before being transferred back on standard diet or on nutrient restriction (NR) diet, water with 7.02 g/L agar only, for 24 hours stress.

### *Ex vivo* CNS cultures and iGluSnFR recordings

Larvae were collected at ∼72h ALH and CNSs dissected in PBS. Concerning the experiment with metabolite, CNSs were kept 2h on PBS only, referred to minimal medium (MM) or with one metabolite. Then, 50 µM EdU was added for an extra hour and the CNSs were fixed with 8% PFA. For isotope labelling experiments, CNSs were incubated in MM for 3 hr with 10 mM of unlabelled Glutamine (control), glutamine-^15^N_2_, glutamine-amide-^15^N_1_, glutamine-amine-^15^N_2_ or glutamate-^13^C_5_. GC-MS quantitation of glutamate excretion into the medium was possible with glutamate-^13^C_5_ but not glutamine-^15^N_2_ as the commercial source of the latter was contaminated with trace amounts of glutamate-^15^N_1_.

For the iGluSnFR.A184A live imaging time-lapse recordings, *ex vivo* CNSs were stabilized in position in an 35 mm x 10 mm petri dish as described^113^ but the tungsten rod was replaced by a thin wire touching the ventral surface and silicone grease was used instead of Blu Tack. After equilibration of the ex vivo CNSs for 2 hr in MM and then the CNSs were viewed with Leica DM6000B microscope under a 10x or 25x water objective. Images were acquired using Leica MM AF 2.2.0 software at 250 ms per frame at a resolution of 256 x 256 pixels using an Orca-Flash 4.0 camera. In time-lapse experiments, at 100 seconds, fresh MM or 20 mM Gln added to a final concentration of 10 mM) and the recording continued for 400 seconds. For iGluSnFR.A184A live imaging at a single timepoint, CNSs were dissected and mounted in PBS. Images were acquired with a 20X objective using a SP8 confocal microscope and LAS X software.

### Tissue staining and confocal microscopy image analysis

All fixation, incubation and wash steps were performed at room temperature in 9-well Pyrex glass plates containing 200 µL solution on a Gyro-Rocker SSL3. Details of antibodies, fluorophores, and concentrations used can be found in the **Table 4**. Larvae CNSs were dissection with sharp tweezers and a tungsten needle in PBS and fixed in 4% (v/v) paraformaldehyde in PBS for 45 min. After 3x PBS washes for 5 min, then permeabilized with 0.2% Triton X-100 in PBS (PBT) for 15 min, next blocked for 30 min in 10% heat-inactivated bovine serum albumin (BSA) in PBS containing 0.2% Triton X-100 (PBT) and subsequently incubated overnight at 4°C with the primary antibody diluted in 10% BSA in PBT. CNSs were then washed in PBT 3x 5min and incubated for 4 hours in the appropriate fluorophore-conjugated secondary antibody diluted in 10% BSA in PBT and then washed in PBT for 3 x 5min. Finally, CNSs were incubated in DAPI (1:2000) in PBT for 15min and washed in PBS for 3 x 5min. Samples were mounted in Vectashield on 21-well diagnostic microscope slides and sealed with glass coverslips (No. 1.5) with nail varnish. Slides were stored at 4°C before imaging. All samples were imaged on either a Leica SP5 upright or a Leica SP8 inverted confocal scanning microscope. Samples were imaged using the same settings (laser power and gain) and a 20X or 40X/1.30 oil-based objectives. The VNC was acquired at a frequency of 400 Hz using 1.5 μm step size z-stacks, starting from the ventral superficial surface until the start of the neuropil.

### Neuroblast proliferation assay

After 24 hours on Fed or NR condition (∼90 hours ALH), larval CNSs were dissected as described above. CNSs were incubated 1 hour with 50 µM 5-ethynyl-2′-deoxyuridine (EdU). Proliferative cells were labelled using the Click-iT™ EdU Cell Proliferation Kit for Imaging, Alexa Fluor™ 555 dye. CNSs were then washed in PBS for 5 min, fixed in 4% PFA in PBS for 45 min, washed in PBT for 3x 5 min, and blocked in 10% BSA in PBT for 30 min. EdU detection was done by incubating the CNSs in the Click-iT reaction cocktail, as described by the manufacturer, for 30min. CNSs were washed 3x 5 min in PBT and processed for Miranda antibody staining and imaging as described above^114^. Neuroblast proliferation is expressed as the proportion of counted Miranda (Mira) expressing NBs that label with EdU in the ventral nerve cord (VNC).

### snRNA-seq

Sample preparation for snRNAseq was based on a previous study^115^. Before starting, all materials were cleaned with ethanol 70% followed by RNAzap, and the solution was kept on ice. Fresh homogenization and resuspension buffers were always prepared. A multi-well glass dish with cold 1x PBS was prepared. The CNS was dissected in one well and then transferred to another well of cold 1x PBS to clean each CNS. The CNS was then transferred using a P1000 tip (coated with BSA) into 500 µL of cold 1x PBS in a nuclease-free 1.5 mL tube on ice (coated with BSA). Samples were frozen at −80°C.

Samples were spun down using a benchtop centrifuge, and the medium was discarded. The CNS was resuspended using a P1000 tip (coated with BSA) in 1 mL homogenization buffer (avoiding bubbles with TritonX-100). A 1 mL homogenized sample was transferred into the 1 mL Dounce. The Dounce Tissue Grinder was heat cleaned at 180°C for 3 hours. Nuclei were released on ice by performing 20 strokes with the loose pestle and 40 strokes with the tight pestle, ensuring that bubbles with Triton-X 100 were avoided. The strainer was first coated with BSA, and then 1 mL of the sample was filtered through a 20 µm cell strainer into a nuclease-free 1.5 mL tube on ice (coated with BSA). Next, the sample was centrifuged for 10 min at 1,000 x g at 4°C. The supernatant was discarded without disturbing the pellet.

The pellet was washed with a P1000 tip (coated with BSA) using 500 µL resuspension buffer. Pipetting was performed > 20 times to completely resuspend the pellet. The sample was again centrifuged for 10 min at 1,000 x g at 4°C, and the supernatant was discarded without disturbing the pellet. The nuclei were resuspended using a P200 tip (coated with BSA) with 150 µL resuspension buffer, pipetting > 20 times to ensure complete resuspension. The sample was kept on ice. Samples were then ready for 10X Genomics. Buffer composition and reagents were listed in **Table 4**.

The concentration and viability of single-cell suspensions were measured using acridine orange (AO) and propidium iodide (PI) with the Luna-FX7 Automatic Cell Counter. Approximately 14,500 cells were loaded onto the Chromium Chip and partitioned into nanoliter-scale droplets using the Chromium X and Chromium GEM-X Single Cell Reagents (Chromium GEM-X Single Cell 3’ v4 Gene Expression User Guide, CG000731). Within each droplet, the cells were lysed, and the RNA was reverse transcribed. All resulting cDNA within a droplet shared the same cell barcode. Illumina-compatible libraries were generated from the cDNA using Chromium GEM-X Single Cell library reagents following the manufacturer’s instructions (Chromium GEM-X Single Cell 3’ v4 Gene Expression User Guide, CG000731). Final libraries were QC’d using the Agilent TapeStation and sequenced using the Illumina NovaSeq X. Sequencing read configuration: 28-10-10-90.

### snRNA-seq data analysis

Libraries were sequenced using the Illumina NextSeq 6000 instrument configured according to 10x recommendations. Gene expression and cell identity were calculated from the 28 base pair (bp) R1 and 90 bp R2 FastQ files using Cell Ranger (8.0.0), implemented by the nf-core^116^ scrnaseq (2.5.1) (https://zenodo.org/records/10554425) Nextflow (24.04.2) pipeline. Unfiltered feature matrices from Cell Ranger were analysed with CellBender (0.3.2)^117^ to remove background signal. Seurat (5.1.0)^118^ objects were created from the filtered CellBender data in R (4.4.1) which were further filtered to remove low quality nuclei using the methods of Fly Cell Atlas^119^. First, doublets were predicted by scDblFinder (1.18.0)^120^ and removed along with nuclei that reported more than 5% mitochondrial reads. Subsequently, the number of UMIs and features were filtered to remove cells outside 3 median absolute deviations (MADs) of the log-transformed median, ensuring that cells provided a minimum of 500 UMIs or 200 features. During cell type annotation, three clusters comprising 896 cells that lacked marker feature expression were identified in the “90h_CNS_5” dataset and removed from the analysis.

Filtered replicate datasets were analysed using Seurat: normalisation, identification of variable features and scaling data used default parameters, and the first 50 principal components were calculated for the PCA reduction. The first 15 (or 18 for 90h_CNS_5) components were used for tSNE and UMAP reductions and neighbour finding. Clusters were identified using FindClusters with default parameters at a range of resolutions. Cell type markers were used to annotate cell clusters at an appropriate resolution; a cluster was subclustered using FindSubCluster if its cells reported markers for multiple cell types. A pooled dataset was created from the replicates using the merge function and reanalysed using the steps outlined above with the first 20 principal components used for dimensionality reduction and neighbour finding. Clusters were defined by following gene markers: neuroblasts/progenitors (*dpn, mira, nab, ase, grh, tll, ovo, eya, cas, sprt, M7BP*), mature larval neurons (*nSyb, Syn, jeb, CG3961, Rdl, para*), immature adult neurons (*pros, nerfin-1, bab1, bab2, ap*), motor neurons (*twit, Proc*), Kenyon cells (*Dop1R2, Rgk1, jdp*), perineurial glia (*daw, vkg, dmGlut, MFS14, scaf*), subperineurial glia (*moody, rost, CG10702, Mdr65*), cortex glia (*Cyp4g15, apolpp, Lip4, ACC*), ensheathing glia (*wrapper, tsl, Oatp26F, CG12880*), astrocyte-like glia (*alrm, CG30197, CG9691, Gat, CG7084*), wrapping glia (*htl, CG42346, fog, Ama, CAP*), and trachea (*verm, pio*).

### GC-MS sample preparation

All solutions/samples were kept on ice (or 4°C when centrifuging) during sample preparation. Metal tubes (2 mL) used for homogenisation were resealed using PTFE O rings (6.07 mm ID x 1.78 mm CS). For larval hemolymph, all samples were collected in a modified VDTS protocol, adapted for GC-MS from a previous published method for NMR^67^. 10 larvae for CW/NR conditions and 5 larvae for the fed condition were washed in saline solution (4.5 g NaCl in 1 L H_2_O) and dried with paper towel before putting into a 60 µL droplet containing 10 nmol norleucine in saline on a lid of the cell culture dish. After 1 minute of contact, 22.5 µL solution in the droplet was taken with a positive displacement syringe and transferred to a Safe-Lock tube (as “pre” sample). Larvae were opened inside the droplet and another 22.5 µL of solution was transferred to another Eppendorf tube (as “post” sample). 100 µL of distilled water was added to both “pre” and “post” samples and then filtered through a 0.22 µm filter unit by centrifuging for 2 minutes at 17 x g to remove hemocytes. Filtered “pre” and “post” samples were transferred to Eppendorf tubes and the following solutions were added: 350 µL chloroform, 196.6 µL methanol and 3.4 µL 1.5 mM ^13^C-3-L-Leucine (in methanol). Samples were vortexed and phase separated by centrifuging for 2 minutes at 17 x g. Separated phases were collected into glass GC vials, for which the upper phase was collected for polar metabolite analysis and stored at −80°C before derivatization, while the lower phase was collected for apolar analysis and stored at −20°C before derivatization. The interphases were kept in the original Eppendorf tubes and stored at −20°C before protein measurement.

For larval CNS analysis, 20 CNSs were dissected in Ringer’s and washed with Ringer’s solution (7.5 g NaCl, 0.35g KCl, 0.21g CaCl_2_ in 1 L H_2_O). 50µL of Ringer’s solution containing tissues were transferred to 2ml metal tubes with 350 µL chloroform, 150 µL methanol, 50 µL 100 µM ^13^C-1-Palmitic acid (in chloroform) and 50 µL 100 µM norleucine. Tissues were homogenised using the Precellys Homogenizer with the standard programme. The solution containing homogenised tissues was transferred to an Eppendorf tube with 150 µL of distilled water added. The Eppendorf tubes were vortexed, phase separated and stored as previously described. At least 5 independent biological replicates were prepared for each condition in all GC-MS experiments.

For *ex vivo* stable isotope tracing experiments, 20 dissected larval CNSs were incubated in 200 µL of ^13^C_5_-glutamine, ^15^N_2_-glutamine or unlabelled glutamine (in Ringer’s solution) for 3 hours. The CNSs were collected and washed in Ringer’s solution and then homogenised as described above (C21:0 is used as internal standard instead of ^13^C-1-Palmitic acid for apolar samples in this experiment). Both polar and apolar phases were transferred to GC vials containing glass inserts and dried in a SpeedVac at 1000 min^-1^ rotation, 10 mbar vacuum and 40°C temperature. 30 µL of methanol was added to the dried sample, which was then re-dried in the SpeedVac, and this was repeated three times to completely dry the samples. Polar phase samples were first derivatised by adding 20 µL of freshly prepared 20 mg/ml methoxyamine hydrochloride in pyridine, vortexing to mix and incubate overnight at room temperature (RT) for complete reaction. The samples were then incubated with 20 µL of BSTFA (+ 1% TMCS) for 1 hour at RT and then run immediately. The apolar phase samples were resuspended in 20 µL of 2:1 chloroform:methanol, vortexing to mix and then derivatised by adding 5 µL of tetramethylammonium hydroxide and then run immediately.

### GC-MS data acquisition and analysis

The GC method was performed using an Agilent 7890B-7000C or 7890B-5977A GC-MS system in EI mode. For polar metabolites, splitless injection of 1μl onto a J&W DB-5ms GC Column with an inlet temperature of 270°C, with helium as the carrier gas. The initial oven temperature was 70°C (2 min), followed by temperature gradients to 295°C at 12.5°C/min, and from 295 to 320°C at 25°C/min with a hold time of 3 min. For FAME analysis of fatty acids, the inlet temperature was 250 °C with an initial oven temperature of 70°C (1 min), followed by temperature gradients to 230 °C at 15 °C/min with a hold time of 2 min, and from 230 to 325 °C at 25 °C/min with a hold time of 3 min. The sample order was randomized before injection. Data was acquired and analysed using MassHunter. Quantification of ^13^C stable isotope data was performed using MANIC, an in-house developed adaptation of the GAVIN package^121^, which also performs correction for natural abundance. ^15^N stable isotope data correction for natural abundance was performed using IsoCorrectoR^122^. Metabolites were identified according to their molecular ion mass and fragmentation patterns in reference to a mixture of commercial standards where possible, details in **Table 5**. Absolute quantifications were carried out only for compounds in reference to commercial standards of known concentration.

### OrbiSIMS sample incubation and preparation

Dissected larval CNS were dissected as described previously and incubated in MM containing either ^13^C ^15^N_5_-deoxyadenosine plus ^15^N-glutamine amide or ^13^C-thymidine plus ^15^N-glutamine amide, for 3 hr. The CNSs were collected, washed in PBS, and fixed in 4% PFA in PBS for 45 min at RT. After fixation, samples were washed in PBS and stained with 13.1 μM Toluidine Blue O for 30 min at RT, followed by a PBS wash. CNSs were then embedded in 4% carboxymethyl cellulose in PBS and flash frozen in a bath of dry ice and 2-methylbutane. Larval CNS transverse sections were cut using a Leica CM3050 S Cryostat set to 10 μm thickness, and −20°C chamber and object temperatures. Tissue slices were thaw mounted onto ITO slides (25 mm x 25 mm) with a resistivity of 70-100 Ω/sq, cleaned via sequential washes in 70:30 acetone:water, 2:1 chloroform:methanol and then hexane, and subsequently vacuum packed and stored at −80°C until analysis.

### OrbiSIMS data acquisition and analysis

CNS section slides were analysed on an OrbiSIMS (HybridSIMS, IONTOF GmbH) instrument at ∼25°C as previously described^79,80^. Analyses were performed using a 20 keV Ar_3500_^+^ quasi-continuous GCIB analysis beam with a spot size of ∼3 μm using a sawtooth raster mode, a current of ∼13 pA, a duty cycle of 10-15%, a cycle length of 200 μs and surface potential of approximately −30 V. The total ion dose for each image was approximately 3.98 x 10^12^ ions cm^-2^ with 1 shot per pixel. Image areas are between 175 x 175 and 200 μm × 200 μm, with a 5 μm pixel size. The images from U-^13^C adenosine samples are the sum of 6 consecutive scans, and ^13^C thymidine samples display the sum of 2 or 3 scans (glutamine ^15^N amide or ^15^N amine) The Orbitrap was operated with a mass resolution of 240,000 @ 200 *m/z* and an injection time of 2961 ms, with the automatic gain control switched off. Mass spectral information was acquired for the mass range m/z 100-350. Putative peak annotations are based on exact mass and isotope distribution analysis, and the assignments are displayed in **Table 5**. The OrbiSIMS instrument was controlled using SurfaceLab, integrating an application programming interface provided by ThermoFisher Scientific. Image analyses were performed using SurfaceLab and Prism. All analysis were performed on the summed signal from all consecutive scans. ROI selections for whole CNS and neuropil regions were performed manually, with delineations based on adenine intensity normalised to total ion count to define the cortex. Images shown correspond to ROI selections of the CNS (outer dotted line) that exclude the surrounding embedding medium. For ratiometric images, the minimum scale value was set at the value corresponding to the natural enrichment of each isotope (values above natural enrichment appear on the colour scale, those at or below this level appear as black).

### Statistical analyses

All statistical analyses were performed in Prism v10 (GraphPad software Inc., San Diego, CA). When using parametric tests (analysis of variance (ANOVA) and *t*-tests), the data were checked for gaussian distribution using either D’Agostino-Pearson omnibus, Anderson-Darling, or Shapiro-Wilk normality tests. The Mann-Whitney test was used for non-parametric tests and the Tukey test for multiple comparisons with ANOVA. Each experiment has n≥10 biological replicates and all experiments were performed at least twice on different days. Significance values: \**p* < 0.05, \*\**p* < 0.01, \*\*\**p* < 0.001, \*\*\*\**p* < 0.0001.

## Supporting information

Extended Video 1: control FRT clones

Extended Video 2: Gs1[1] FRT clones

## Extended Data Figure Legends

**Extended Data Figure 1:**
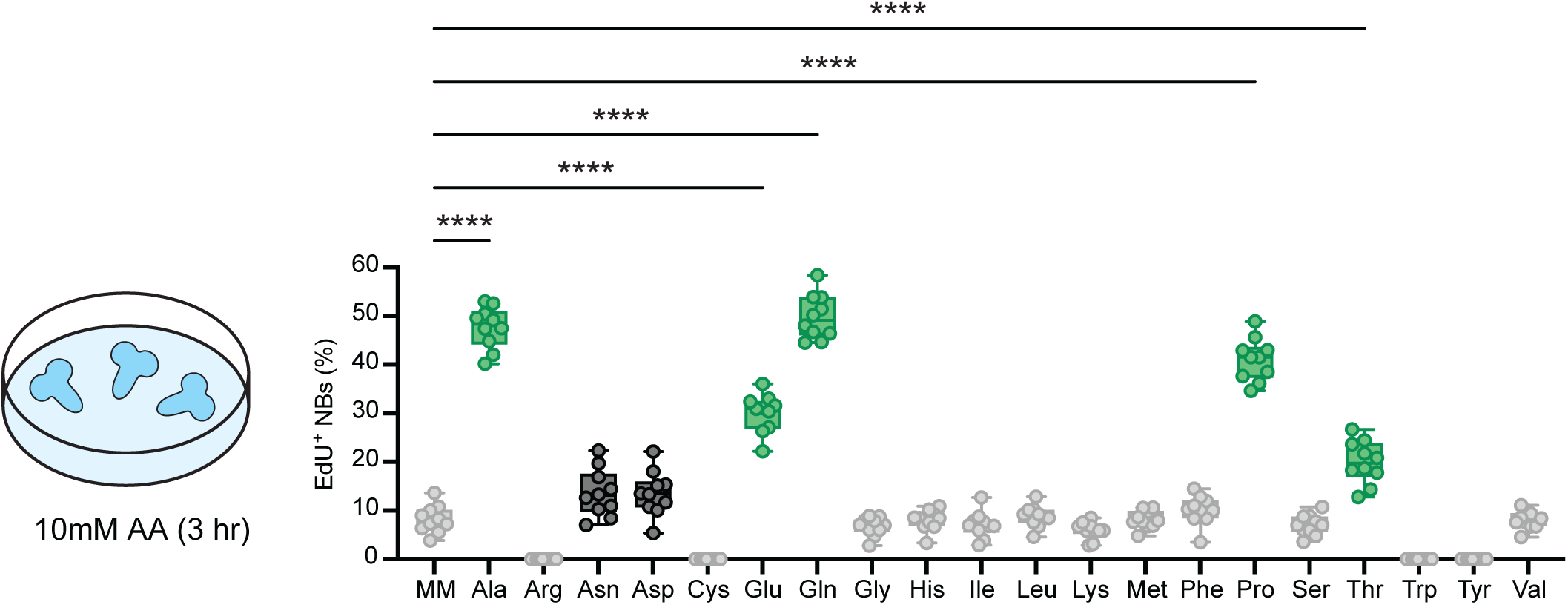
High concentrations of Ala, Gln, Glu, Pro or Thr support *ex vivo* neuroblast proliferation. **a**, CNSs were cultured for 3 hr with 10 mM AA. Ala, Glu, Gln, Pro, and Thr in MM supported sufficient neuroblast proliferation (green). Incubation with 10 mM Asn and Asp slightly stimulated neuroblast proliferation (black). One-way ANOVA with multiple comparisons was performed: \**p* < 0.05 for black boxes; \*\*\*\**p* < 0.0001 for green boxes.

**Extended Data Figure 2:**
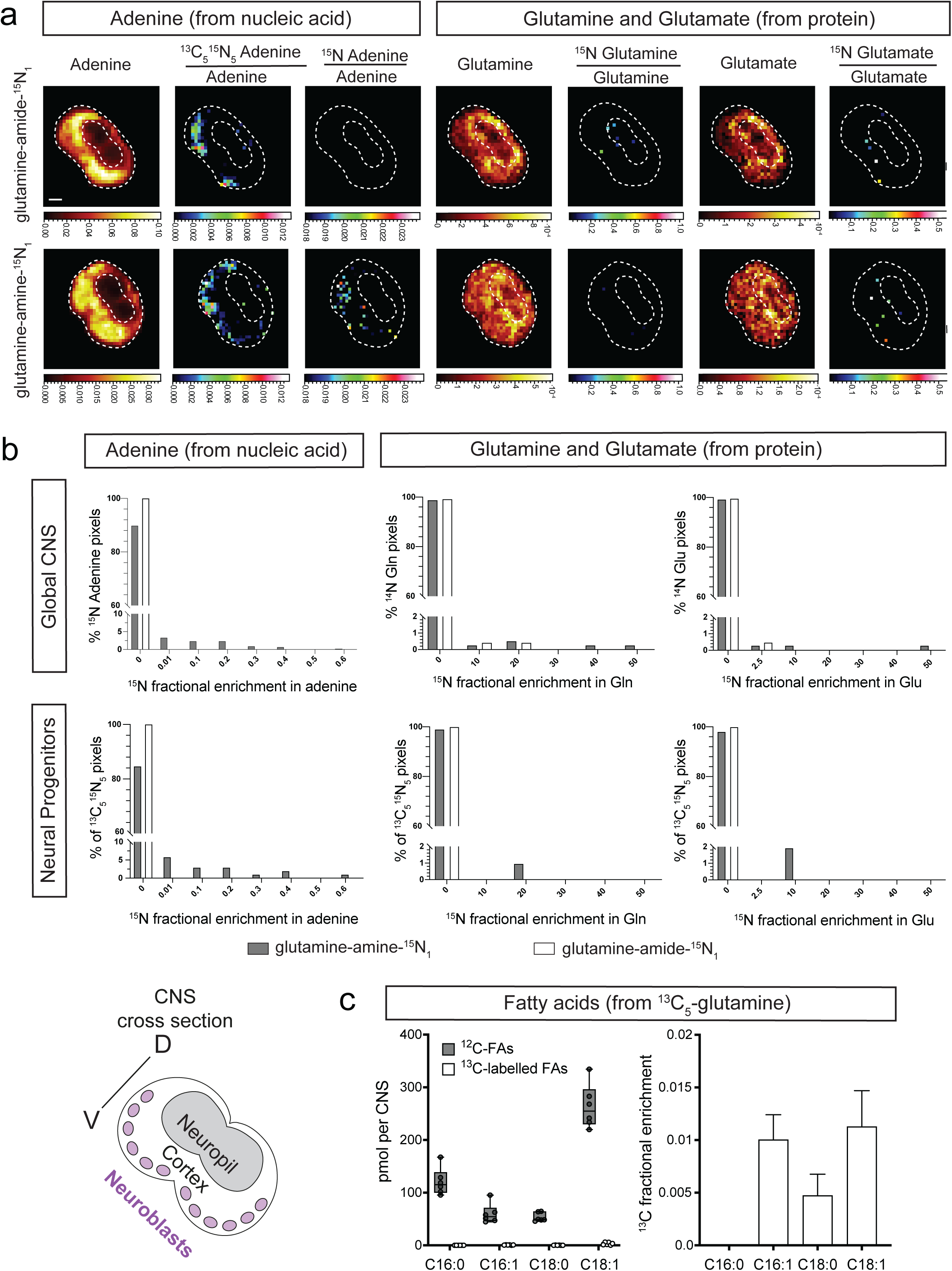
Global CNS and progenitor-specific isotope tracing of glutamine nitrogen. **a**, OrbiSIMS images of cross-sections of CNSs following *ex vivo* incubation for 3 hr in minimal medium supplemented with ^13^C_10_^15^N_5_-deoxyadenosine and Gln-amide-^15^N_1_ (top row) or Gln-amine-^15^N_1_ (bottom row). Pixels with ^13^C_5_^15^N_5_ adenine signals indicate dividing neural progenitors, located in the cortex of the CNS. Only low incorporation of ^15^N from Gln-amide-^15^N_1_ or Gln-amine-^15^N_1_ into the adenine of DNA or the glutamine or glutamate (detected as glutamic acid) of protein was detected within neural progenitors or more globally throughout the CNS. Labelled ions (^13^C_5_^15^N_5_ adenine, ^15^N adenine, ^15^N glutamine and ^15^N glutamate) are displayed as ratios over unlabelled adenine, glutamine, or glutamate. The minimum scale values represent natural isotope abundance. Unlabelled ions (adenine, glutamine, and glutamate) are displayed as ratios over the total ion count. Cortex and neuropil compartments of the CNS (white dotted lines) were delineated using the adenine ion. For ^15^N adenine, enrichment is displayed above natural abundance but the uncorrected total ion intensities are high, ranging between 2.021×10^5^ - 2.295×10^5^. **b**, Quantifications of OrbiSIMS images in (a). Neural-progenitor specific and global CNS (cortex) fractional enrichments of ^15^N into the adenine of DNA and the glutamine or glutamate of protein are shown as pixel frequency distributions. The y-axis shows the percentage of double-positive pixels (y axis) and the ^15^N fractional enrichments (above natural abundance) are binned along the x-axis. **c**, Glutamine carbon incorporation into the major fatty acids of CNSs (schematic, left) incubated in minimal medium supplemented with ^13^C_5_-glutamine. Graphs show GC-MS quantifications of the amounts (centre) and the ^13^C fractional enrichments (right) of the major fatty acids (C16:0, C16:1, C18:0, C18:1).

**Extended Data Figure 3:**
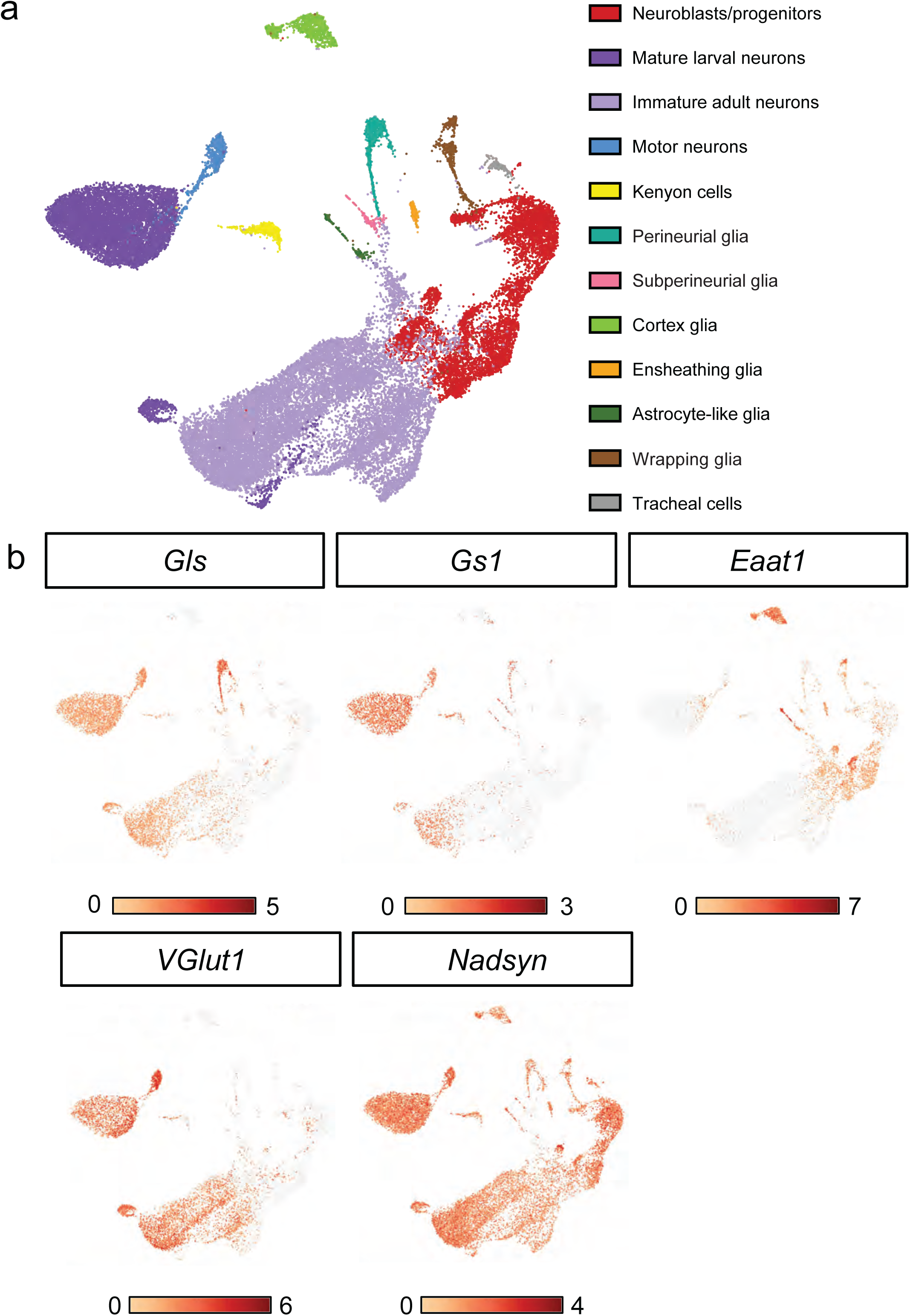
snRNAseq of the larval CNS at 90 hr ALH. **a**, A UMAP of five single-nuclear transcriptomic replicates of 90h ALH CNSs showing the two-dimensional projection of 12 major cell types (27,794 cells). The count of each cell type: neuroblasts/progenitors (4,161), mature larval neurons (7,426), immature adult neurons (12,326), motor neurons (651), Kenyon cells (541), perineurial glia (664), subperineurial glia (156), cortex glia (933), ensheathing glia (233), astrocyte-like glia (197), wrapping glia (516), and trachea (200). **b**, Feature plots showing normalised expression of *Gls*, *Gs1*, *Eaat1*, *VGlut1*, and *Nadsyn* expression in the CNS.

**Extended Data Figure 4:**
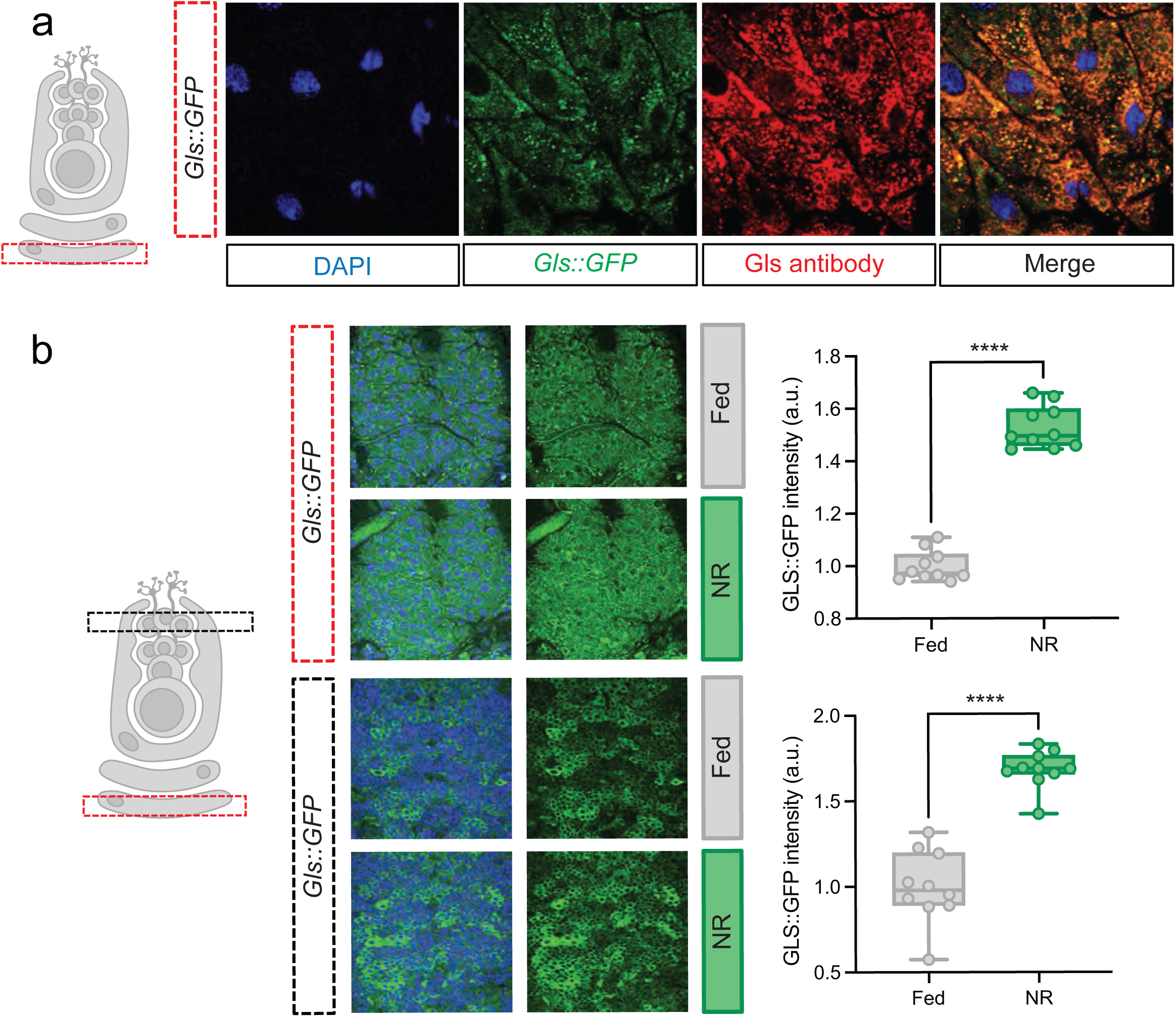
Gls is induced during NR in perineurial glia and mature larval neurons. **a**, Confocal image of Gls::GFP expression and Gls antibody detection in the BBB, showing complete colocalization. **b**, Quantification of Gls::GFP in the BBB (red) and MLN (black) regions under fed and NR conditions, revealing a significant increase in GFP signal under NR.

**Extended Data Figure 5:**
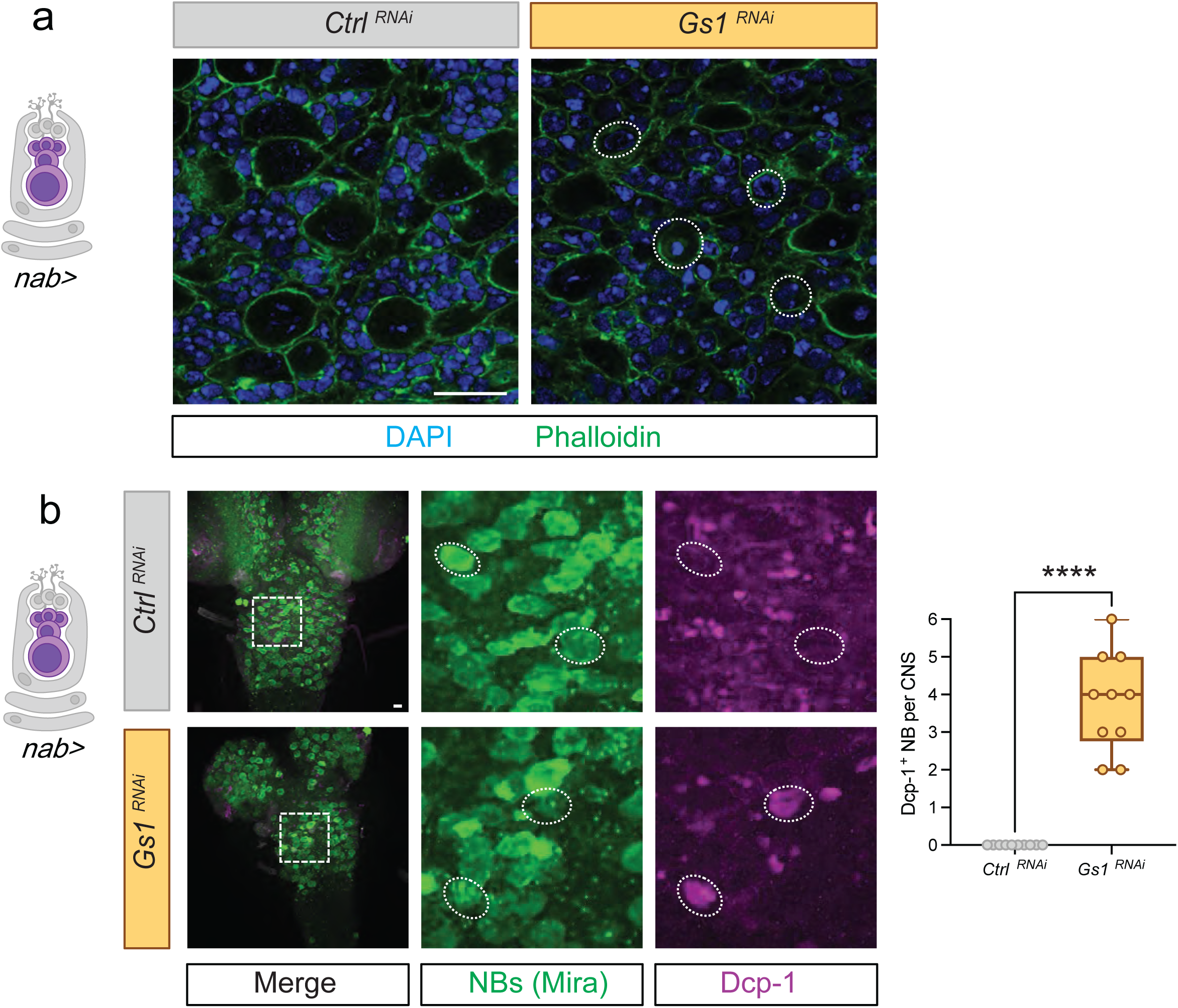
Some *Gs1*-deficient neuroblasts undergo apoptosis. **a**, Apoptotic cells were detected in CNSs using Dcp-1 antibody. In control CNSs, only some neurons were positive for Dcp-1 staining, whereas Gs1 inhibition in NBs resulted in neural stem cells positive for Dcp-1. At 90h ALH, approximately four NBs per CNS were apoptotic compared to none in control CNSs. **b**, CNSs with Gs1 inhibition in NBs exhibited abnormal nuclear structures. Some round cells with compact DAPI staining appeared larger than neurons but smaller than NBs, which were absent in control CNSs. Neuroblasts were labelled with Mira antibody (green), apoptosis with Dcp-1 (purple), neurons with DAPI (blue), and actin was visualized with phalloidin (green). Scale bar = 10 µm. Unpaired t-tests were performed: \*\*\*\**p* < 0.0001.

**Extended Data Figure 6:**
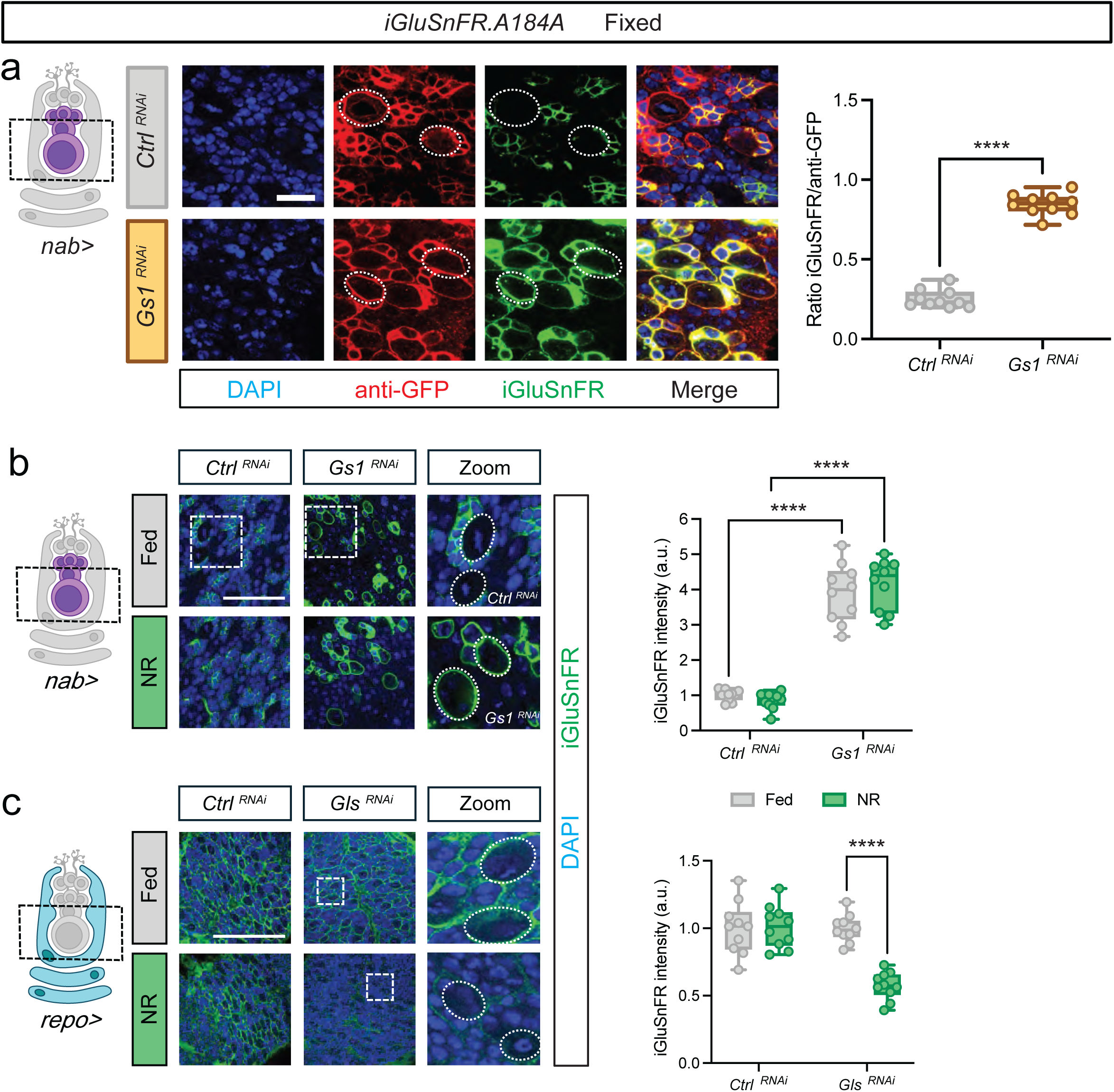
Glial Gls promotes Glu accumulation in cortex glial chambers. **a**, To confirm that Gs1 inhibition in NBs does not alter iGluSnFR expression, an anti-GFP antibody was used. The ratio of GFP from iGluSnFR (green) to anti-GFP (red) remained higher under Gs1 inhibition without variation in the red signal between control and Gs1 inhibition CNSs. **b**, CNSs expressing iGluSnFR under control (mCherry RNAi) or Gs1 RNAi in NBs. CNSs with Gs1 RNAi exhibited an increased iGluSnFR signal compared to controls. **c**, Inhibition of Gls in glial cells using repo-Gal4 led to significant loss of iGluSnFR signal under NR but not fed conditions. Neurons were labelled with DAPI (blue), and iGluSnFR was visualized with GFP (green). Scale bar = 10 µm for (a) and 50 µm for (b, c). Two-way ANOVA with multiple comparisons was performed: \*\*\*\**p* < 0.0001.

**Extended Data Figure 7:**
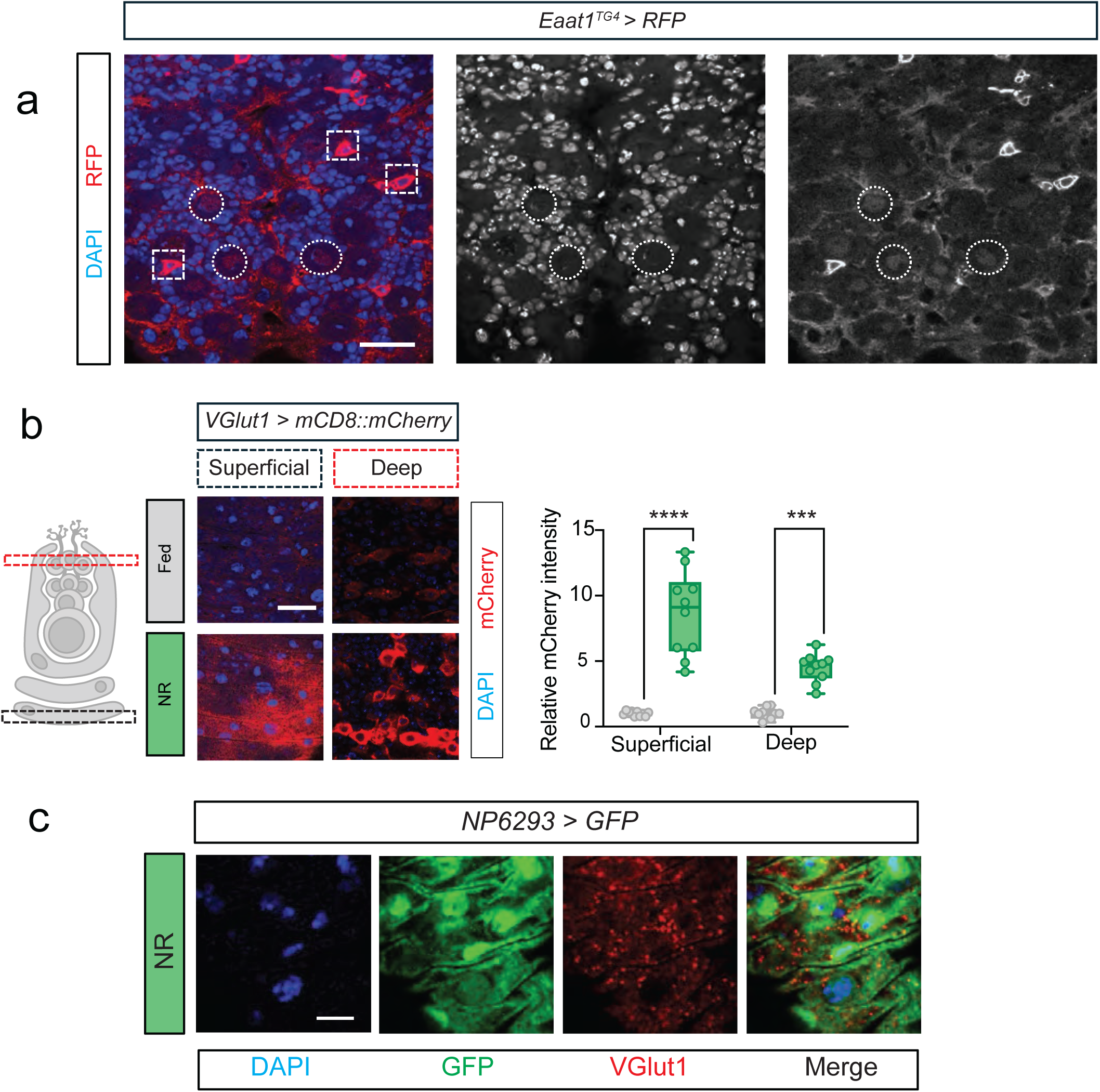
Neuroblasts express Eaat1 and perineurial glia upregulate VGlut1 during NR. **a**, VGlut1 expression was observed using a VGlut1 driver coupled with UAS-mCD8::mCherry. An increase in mCherry signal was detected under NR conditions at the BBB and in mature larval neurons. **b**, To confirm VGlut1 localization in the BBB, a perineural glia driver (NP6293-Gal4) expressing UAS-EGFP was used alongside a VGlut1 antibody. Colocalization of VGlut1 antibody (red) with perineural glial cells (green) was observed. c, Eaat1-expressing cells were visualized using a Trojan-Gal4 line expressing UAS-EGFP, showing a pattern similar to cortex glial cells, consistent with nuclear scRNAseq data. Scale bar = 50 µm for (a) and 10 µm for (b, c). Two-way ANOVA with multiple comparisons was performed: \*\*\*\**p* < 0.0001.

**Extended Data Figure 8:**
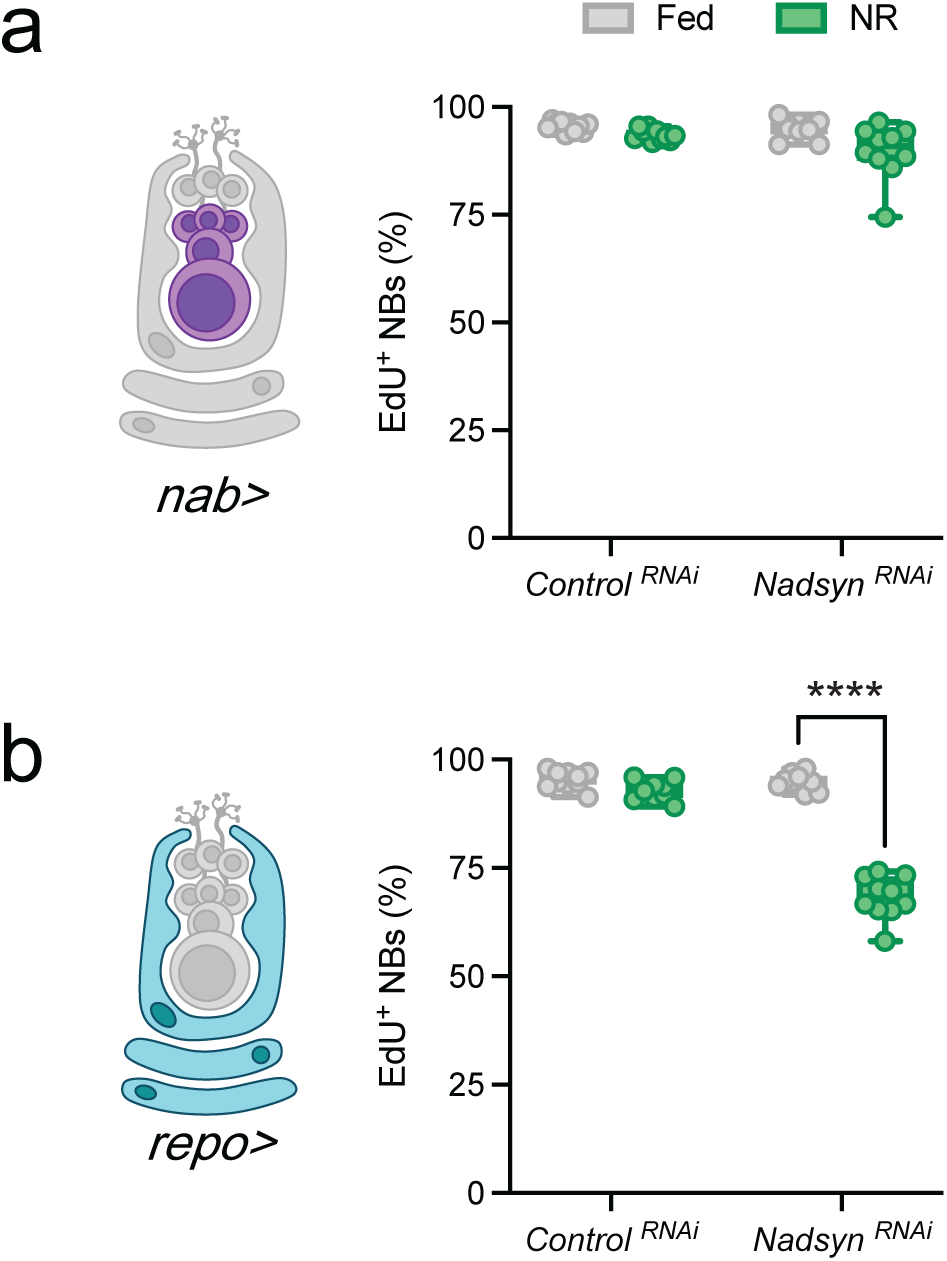
Glial Nadsyn is required for neuroblast proliferation during NR. **a**, Nadsyn inhibition via RNAi in NBs (nab-Gal4) did not affect neural stem cell proliferation under either fed or NR conditions. **b**, Nadsyn inhibition via RNAi in glial cells (repo-Gal4) resulted in reduced NB proliferation under NR conditions, similar to Gls RNAi inhibition in glial cells. Two-way ANOVA with multiple comparisons was performed: \*\*\*\**p* < 0.0001.

## Acknowledgements

We acknowledge Brian McCabe for antibodies and Jonathan Marvin and Loren Looger for iGluSnFR discussions. Fly stocks were also obtained from the Bloomington Drosophila Stock Center (NIH P40OD018537), the Vienna Drosophila Research Centre, and the Kyoto Drosophila Genetic Resource. We thank Vanessa Nunes, James Ellis and the Crick Metabolomics STP and also the Crick Genomics and Bioinformatics and Biostatistics STPs. We also thank Dimitrios Anastasiou, Andrew Bailey, Patricia Serpente, Ying Zhang, Sebastian Sorge, and Elisabeth Kamper for advice and comments on the manuscript. CNS and petri dish images were made using BioRender. This work was supported by an Investigator Award (104566) from the Wellcome Trust, a research grant from NC3Rs (NC/V001272/1) that supported A.F., and by funding from the Francis Crick Institute, which receives its core funding from Cancer Research UK (FC001088), the UK Medical Research Council (FC001088) and the Wellcome Trust (FC001088). For the purpose of Open Access, the author has applied a CC BY public copyright licence to any Author Accepted Manuscript version arising from this submission.

## Author Contributions

Conceptualization, A.F. and A.P.G.; Formal analysis, A.F., Y.J., C.L.N. and J.I.M.; Data curation, C.L.N. and C.B.; Funding acquisition, I.S.G. and A.P.G.; Investigation, A.F., Y.J., C.L.N., V.G and G.M. Methodology, A.F. and A.P.G.; Project administration, A.F. and A.P.G.; Resources, I.S.G. and A.P.G.; Software, C.B., Supervision, A.P.G.; Validation, A.F.; Visualization, A.F. and A.P.G.; Writing - original draft, A.F. and A.P.G.; Writing - review & editing, A.F., Y.J., C.L.N., V.G., G.M., Y.G., C.B., J.I.M., I.S.G., and A.P.G.

## Data Availability

All relevant data can be found within the article and all GC-MS and OrbiSIMS data will be deposited on MetaboLights.

