## Supplementary material for "An intercellular metabolic relay for brain sparing in *Drosophila*": Tables 1,2,3,4 and 5

**Table 1: AA concentration (related to Fig. 1)**

| **Hemolymph** | | | | |
|---|---|---|---|---|
| **Amino Acid** | **[Fed] (mM)** | **+/- SD** | **[NR] (mM)** | **+/- SD** |
| Alanine | 9.50 | 0.72 | 3.41 | 1.15 |
| Arginine | nd | nd | nd | nd |
| Asparagine | 2.40 | 0.54 | 0.62 | 0.35 |
| Aspartate | 0.45 | 0.13 | 0.29 | 0.16 |
| Cysteine | 0.05 | 0.01 | 0.07 | 0.01 |
| Glutamate | 0.07 | 0.02 | 0.07 | 0.04 |
| Glutamine | 4.31 | 3.19 | 1.46 | 1.31 |
| Glycine | 5.54 | 0.66 | 2.95 | 0.86 |
| Histidine | 2.86 | 0.52 | 3.99 | 0.91 |
| Isoleucine | 1.26 | 0.12 | 0.52 | 0.33 |
| Leucine | 1.98 | 0.19 | 0.46 | 0.27 |
| Lysine | 8.95 | 1.20 | 6.00 | 0.89 |
| Methionine | 0.56 | 0.10 | 0.17 | 0.11 |
| Phenylalanine | 0.54 | 0.07 | 0.13 | 0.06 |
| Proline | 7.30 | 0.74 | 1.88 | 0.47 |
| Serine | 3.56 | 0.31 | 1.45 | 0.39 |
| Threonine | 5.74 | 0.55 | 1.14 | 0.36 |
| Tryptophan | 0.05 | 0.04 | 0.01 | 0.00 |
| Tyrosine | 3.04 | 0.39 | 0.96 | 0.71 |
| Valine | 2.77 | 0.24 | 0.91 | 0.48 |

| **CNS** | | | | |
|---|---|---|---|---|
| **Amino Acid** | **Fed (pmol)** | **+/- SD** | **NR (pmol)** | **+/- SD** |
| Alanine | 5.13 | 1.64 | 2.43 | 0.76 |
| Arginine | 0.00 | 0.00 | 0.00 | 0.00 |
| Asparagine | 0.05 | 0.02 | 0.00 | 0.02 |
| Aspartate | 0.70 | 0.33 | 0.38 | 0.28 |
| Cysteine | 0.00 | 0.00 | 0.00 | 0.00 |
| Glutamate | 1.04 | 0.41 | 0.54 | 0.29 |
| Glutamine | 0.05 | 0.03 | 0.00 | 0.03 |
| Glycine | 1.31 | 0.41 | 0.29 | 0.41 |
| Histidine | 0.00 | 0.00 | 0.00 | 0.00 |
| Isoleucine | 0.31 | 0.09 | 0.06 | 0.09 |
| Leucine | 0.21 | 0.08 | 0.04 | 0.03 |
| Lysine | 0.72 | 0.23 | 0.12 | 0.25 |
| Methionine | 0.00 | 0.00 | 0.00 | 0.00 |
| Phenylalanine | 0.04 | 0.01 | 0.01 | 0.01 |
| Proline | 2.92 | 0.86 | 0.68 | 0.78 |
| Serine | 0.51 | 0.16 | 0.11 | 0.15 |
| Threonine | 1.57 | 0.40 | 0.15 | 0.30 |
| Tryptophan | 0.00 | 0.00 | 0.00 | 0.00 |
| Tyrosine | 0.66 | 0.12 | 0.14 | 0.13 |
| Valine | 0.82 | 0.24 | 0.13 | 0.19 |

### Table 2: Isotope enrichments (related to Fig. 2)

**Absolute amount quantification from the $^{13}$C5-Gln experiment**

**Corrected amount of metabolite in CNS (pmol per CNS)**

| | Glutamine | | | | | | |
|---|---|---|---|---|---|---|---|
| | M+0 | M+1 | M+2 | M+3 | M+4 | M+5 | Summed pmol |
| 1 | 0.00 | 0.00 | 0.00 | 0.00 | 0.00 | 271.72 | 271.72 |
| 2 | 0.00 | 0.00 | 0.00 | 0.00 | 0.00 | 3955.77 | 3955.77 |
| 3 | 0.00 | 0.00 | 0.00 | 0.00 | 0.00 | 2913.61 | 2913.61 |
| 4 | 0.00 | 0.00 | 0.00 | 0.00 | 0.00 | 2388.43 | 2388.43 |
| 5 | 0.00 | 0.00 | 0.00 | 0.00 | 0.00 | 2676.73 | 2676.73 |
| 6 | 0.00 | 0.00 | 0.00 | 0.00 | 0.00 | 6018.38 | 6018.38 |
| Average | 0.00 | 0.00 | 0.00 | 0.00 | 0.00 | 3037.44 | 3037.44 |

| | Glutamate* | | | | | | |
|---|---|---|---|---|---|---|---|
| | M+0 | M+1 | M+2 | M+3 | M+4 | M+5 | Summed pmol |
| 1 | 6.42 | 0.46 | 1.88 | 0.00 | 9.25 | | 18.02 |
| 2 | 14.37 | 1.69 | 5.73 | 0.00 | 29.27 | | 51.07 |
| 3 | 10.91 | 1.22 | 3.83 | 0.07 | 20.06 | | 36.09 |
| 4 | 7.20 | 1.26 | 4.13 | 0.07 | 20.40 | | 33.06 |
| 5 | 7.32 | 1.21 | 3.40 | 0.11 | 17.52 | | 29.56 |
| 6 | 7.54 | 1.83 | 4.76 | 0.07 | 25.41 | | 39.61 |
| Average | 8.96 | 1.28 | 3.96 | 0.05 | 20.32 | | 34.57 |

| | GABA | | | | | | |
|---|---|---|---|---|---|---|---|
| | M+0 | M+1 | M+2 | M+3 | M+4 | M+5 | Summed pmol |
| 1 | 0.87 | 0.00 | 0.90 | 0.00 | 14.24 | | 16.01 |
| 2 | 0.68 | 0.00 | 1.08 | 0.00 | 19.75 | | 21.51 |
| 3 | 0.39 | 0.00 | 0.87 | 0.00 | 17.87 | | 19.14 |
| 4 | 0.38 | 0.00 | 0.77 | 0.00 | 20.28 | | 21.43 |
| 5 | 0.81 | 0.00 | 1.25 | 0.00 | 19.87 | | 21.93 |
| 6 | 0.09 | 0.27 | 1.55 | 0.00 | 25.33 | | 27.24 |
| Average | 0.54 | 0.05 | 1.07 | 0.00 | 19.56 | | 21.21 |

| | Alanine | | | | | | |
|---|---|---|---|---|---|---|---|
| | M+0 | M+1 | M+2 | M+3 | M+4 | M+5 | Summed pmol |
| 1 | 7.36 | 1.09 | 6.02 | | | | 14.48 |
| 2 | 19.25 | 1.81 | 10.37 | | | | 31.43 |
| 3 | 10.82 | 1.43 | 7.37 | | | | 19.62 |
| 4 | 10.42 | 1.28 | 7.18 | | | | 18.87 |
| 5 | 9.89 | 1.12 | 6.54 | | | | 17.56 |
| 6 | 22.47 | 2.33 | 12.73 | | | | 37.53 |
| Average | 13.37 | 1.51 | 8.37 | | | | 23.25 |

| | Pyruvate | | | | | | |
|---|---|---|---|---|---|---|---|
| | M+0 | M+1 | M+2 | M+3 | M+4 | M+5 | Summed pmol |
| 1 | 3.03 | 0.12 | 0.12 | 0.79 | | | 4.06 |
| 2 | 3.00 | 0.11 | 0.00 | 0.88 | | | 3.98 |
| 3 | 3.30 | 0.11 | 0.18 | 1.21 | | | 4.80 |
| 4 | 3.50 | 0.17 | 0.06 | 0.92 | | | 4.66 |
| 5 | 0.85 | 0.12 | 0.12 | 0.88 | | | 1.97 |
| 6 | 4.22 | 0.19 | 0.34 | 2.06 | | | 6.82 |
| Average | 2.98 | 0.14 | 0.14 | 1.12 | | | 4.38 |

| | Succinate | | | | | | |
|---|---|---|---|---|---|---|---|
| | M+0 | M+1 | M+2 | M+3 | M+4 | M+5 | Summed pmol |
| 1 | 1.18 | 0.00 | 0.50 | 0.00 | 1.95 | | 3.63 |
| 2 | 1.87 | 0.01 | 0.52 | 0.00 | 2.91 | | 5.30 |
| 3 | 1.04 | 0.00 | 0.49 | 0.00 | 2.32 | | 3.84 |
| 4 | 1.72 | 0.00 | 0.54 | 0.00 | 2.57 | | 4.83 |
| 5 | 0.83 | 0.00 | 0.34 | 0.00 | 2.03 | | 3.20 |
| 6 | 1.08 | 0.06 | 0.38 | 0.00 | 2.95 | | 4.46 |
| Average | 1.29 | 0.01 | 0.46 | 0.00 | 2.45 | | 4.21 |

| | Fumarate | | | | | | |
|---|---|---|---|---|---|---|---|
| | M+0 | M+1 | M+2 | M+3 | M+4 | M+5 | Summed pmol |
| 1 | 0.28 | 0.04 | 0.08 | 0.03 | 0.34 | | 0.77 |
| 2 | 0.63 | 0.05 | 0.08 | 0.01 | 0.45 | | 1.22 |
| 3 | 0.35 | 0.04 | 0.11 | 0.02 | 0.52 | | 1.05 |
| 4 | 0.21 | 0.03 | 0.10 | 0.00 | 0.46 | | 0.80 |
| 5 | 0.21 | 0.06 | 0.10 | 0.02 | 0.48 | | 0.87 |
| 6 | 0.30 | 0.07 | 0.11 | 0.02 | 0.64 | | 1.14 |
| Average | 0.33 | 0.05 | 0.10 | 0.02 | 0.48 | | 0.98 |

| | Lactate | | | | | | |
|---|---|---|---|---|---|---|---|
| | M+0 | M+1 | M+2 | M+3 | M+4 | M+5 | Summed pmol |
| 1 | 7.43 | 0.16 | 0.94 | | | | 8.52 |
| 2 | 15.28 | 0.08 | 1.55 | | | | 16.92 |
| 3 | 13.22 | 0.19 | 0.53 | | | | 13.94 |
| 4 | 7.36 | 0.09 | 1.11 | | | | 8.57 |
| 5 | 0.00 | 0.00 | 0.68 | | | | 0.68 |
| 6 | 7.62 | 0.54 | 2.88 | | | | 11.04 |
| Average | 8.49 | 0.18 | 1.28 | | | | 9.95 |

**Fraction labelling from the $^{13}$C5-Gln experiment**

**Corrected labelling fraction of metabolite in CNS**

| | Glutamine | | | | | | |
|---|---|---|---|---|---|---|---|
| | M+0 | M+1 | M+2 | M+3 | M+4 | M+5 | Mean enrichment |
| 1 | 0.00 | 0.00 | 0.00 | 0.00 | 0.00 | 1.00 | 1.00 |
| 2 | 0.00 | 0.00 | 0.00 | 0.00 | 0.00 | 1.00 | 1.00 |
| 3 | 0.00 | 0.00 | 0.00 | 0.00 | 0.00 | 1.00 | 1.00 |
| 4 | 0.00 | 0.00 | 0.00 | 0.00 | 0.00 | 1.00 | 1.00 |
| 5 | 0.00 | 0.00 | 0.00 | 0.00 | 0.00 | 1.00 | 1.00 |
| 6 | 0.00 | 0.00 | 0.00 | 0.00 | 0.00 | 1.00 | 1.00 |
| Average | 0.00 | 0.00 | 0.00 | 0.00 | 0.00 | 1.00 | 1.00 |

| | Glutamate* | | | | | | |
|---|---|---|---|---|---|---|---|
| | M+0 | M+1 | M+2 | M+3 | M+4 | M+5 | Mean enrichment |
| 1 | 0.36 | 0.03 | 0.10 | 0.00 | 0.51 | | 0.57 |
| 2 | 0.28 | 0.03 | 0.11 | 0.00 | 0.57 | | 0.64 |
| 3 | 0.30 | 0.03 | 0.11 | 0.00 | 0.56 | | 0.62 |
| 4 | 0.22 | 0.04 | 0.13 | 0.00 | 0.62 | | 0.69 |
| 5 | 0.25 | 0.04 | 0.12 | 0.00 | 0.59 | | 0.66 |
| 6 | 0.19 | 0.05 | 0.12 | 0.00 | 0.64 | | 0.71 |
| Average | 0.27 | 0.04 | 0.11 | 0.00 | 0.58 | | 0.65 |

| | GABA | | | | | | |
|---|---|---|---|---|---|---|---|
| | M+0 | M+1 | M+2 | M+3 | M+4 | M+5 | Mean enrichment |
| 1 | 0.05 | 0.00 | 0.06 | 0.00 | 0.89 | | 0.92 |
| 2 | 0.03 | 0.00 | 0.05 | 0.00 | 0.92 | | 0.94 |
| 3 | 0.02 | 0.00 | 0.05 | 0.00 | 0.93 | | 0.96 |
| 4 | 0.02 | 0.00 | 0.04 | 0.00 | 0.95 | | 0.96 |
| 5 | 0.04 | 0.00 | 0.06 | 0.00 | 0.91 | | 0.93 |
| 6 | 0.00 | 0.01 | 0.06 | 0.00 | 0.93 | | 0.96 |
| Average | 0.03 | 0.00 | 0.05 | 0.00 | 0.92 | | 0.95 |

| | Alanine | | | | | | |
|---|---|---|---|---|---|---|---|
| | M+0 | M+1 | M+2 | M+3 | M+4 | M+5 | Mean enrichment |
| 1 | 0.51 | 0.08 | 0.42 | | | | 0.45 |
| 2 | 0.61 | 0.06 | 0.33 | | | | 0.36 |
| 3 | 0.55 | 0.07 | 0.38 | | | | 0.41 |
| 4 | 0.55 | 0.07 | 0.38 | | | | 0.41 |
| 5 | 0.56 | 0.06 | 0.37 | | | | 0.40 |
| 6 | 0.60 | 0.06 | 0.34 | | | | 0.37 |
| Average | 0.56 | 0.07 | 0.37 | | | | 0.40 |

| | Pyruvate | | | | | | |
|---|---|---|---|---|---|---|---|
| | M+0 | M+1 | M+2 | M+3 | M+4 | M+5 | Mean enrichment |
| 1 | 0.75 | 0.03 | 0.03 | 0.19 | | | 0.22 |
| 2 | 0.75 | 0.03 | 0.00 | 0.22 | | | 0.23 |
| 3 | 0.69 | 0.02 | 0.04 | 0.25 | | | 0.29 |
| 4 | 0.75 | 0.04 | 0.01 | 0.20 | | | 0.22 |
| 5 | 0.43 | 0.06 | 0.06 | 0.45 | | | 0.51 |
| 6 | 0.62 | 0.03 | 0.05 | 0.30 | | | 0.34 |
| Average | 0.66 | 0.03 | 0.03 | 0.27 | | | 0.30 |

| | Succinate | | | | | | |
|---|---|---|---|---|---|---|---|
| | M+0 | M+1 | M+2 | M+3 | M+4 | M+5 | Mean enrichment |
| 1 | 0.32 | 0.00 | 0.14 | 0.00 | 0.54 | | 0.61 |
| 2 | 0.35 | 0.00 | 0.10 | 0.00 | 0.55 | | 0.60 |
| 3 | 0.27 | 0.00 | 0.13 | 0.00 | 0.60 | | 0.67 |
| 4 | 0.36 | 0.00 | 0.11 | 0.00 | 0.53 | | 0.59 |
| 5 | 0.26 | 0.00 | 0.11 | 0.00 | 0.63 | | 0.69 |
| 6 | 0.24 | 0.01 | 0.09 | 0.00 | 0.66 | | 0.71 |
| Average | 0.30 | 0.00 | 0.11 | 0.00 | 0.59 | | 0.64 |

| | Fumarate | | | | | | |
|---|---|---|---|---|---|---|---|
| | M+0 | M+1 | M+2 | M+3 | M+4 | M+5 | Mean enrichment |
| 1 | 0.36 | 0.05 | 0.10 | 0.04 | 0.45 | | 0.54 |
| 2 | 0.52 | 0.04 | 0.07 | 0.01 | 0.37 | | 0.42 |
| 3 | 0.34 | 0.04 | 0.11 | 0.02 | 0.50 | | 0.58 |
| 4 | 0.26 | 0.04 | 0.12 | 0.00 | 0.58 | | 0.65 |
| 5 | 0.24 | 0.07 | 0.12 | 0.02 | 0.55 | | 0.65 |
| 6 | 0.26 | 0.06 | 0.09 | 0.02 | 0.56 | | 0.64 |
| Average | 0.33 | 0.05 | 0.10 | 0.02 | 0.50 | | 0.58 |

| | Lactate | | | | | | |
|---|---|---|---|---|---|---|---|
| | M+0 | M+1 | M+2 | M+3 | M+4 | M+5 | Mean enrichment |
| 1 | 0.87 | 0.02 | 0.11 | | | | 0.12 |
| 2 | 0.90 | 0.01 | 0.09 | | | | 0.09 |
| 3 | 0.95 | 0.01 | 0.04 | | | | 0.05 |
| 4 | 0.86 | 0.01 | 0.13 | | | | 0.14 |
| 5 | 0.00 | 0.00 | 1.00 | | | | 1.00 |
| 6 | 0.69 | 0.05 | 0.26 | | | | 0.29 |
| Average | 0.71 | 0.02 | 0.27 | | | | 0.28 |

**Absolute amount quantification from the $^{15}$N2-Gln experiment**

**Corrected amount of metabolite in CNS (pmol per CNS)**

| | Glutamine | | | | Glutamate | | | |
|---|---|---|---|---|---|---|---|---|
| | M+0 | M+1 | M+2 | Summed pmol | M+0 | M+1 | M+2 | Summed pmol |
| 1 | 158.06 | 0.00 | 13740.85 | 13898.91 | 23.68 | 206.71 | | 230.40 |
| 2 | 127.13 | 0.00 | 10317.11 | 10444.24 | 3.56 | 139.40 | | 142.96 |
| 3 | 158.23 | 0.00 | 14195.92 | 14354.15 | 0.00 | 226.40 | | 226.40 |
| 4 | 87.08 | 0.00 | 9702.06 | 9789.14 | 10.61 | 155.71 | | 166.32 |
| 5 | 110.74 | 0.00 | 9869.18 | 9979.92 | 0.00 | 161.59 | | 161.59 |
| 6 | 476.96 | 0.00 | 29600.18 | 30077.14 | 0.00 | 466.52 | | 466.52 |
| Average | 186.37 | 0.00 | 14570.88 | 14757.25 | 6.31 | 226.06 | | 232.37 |

| | GABA | | | | Alanine | | | | Aspartate | | | |
|---|---|---|---|---|---|---|---|---|---|---|---|---|
| | M+0 | M+1 | M+2 | Summed pmol | M+0 | M+1 | M+2 | Summed pmol | M+0 | M+1 | M+2 | Summed pmol |
| 1 | 2.25 | 44.02 | | 46.28 | 6.05 | 28.85 | | 34.90 | 20.13 | 202.30 | | 222.42 |
| 2 | 3.25 | 34.82 | | 38.07 | 7.49 | 23.46 | | 30.94 | 11.96 | 130.21 | | 142.17 |
| 3 | 2.10 | 37.32 | | 39.42 | 8.29 | 31.18 | | 39.47 | 34.21 | 157.94 | | 192.15 |
| 4 | 4.40 | 39.29 | | 43.69 | 5.56 | 21.61 | | 27.18 | 30.63 | 204.80 | | 235.43 |
| 5 | 2.88 | 41.32 | | 44.20 | 5.83 | 28.11 | | 33.94 | 47.44 | 149.51 | | 196.94 |
| 6 | 0.00 | 46.06 | | 46.06 | 18.96 | 63.60 | | 82.56 | 46.45 | 233.24 | | 279.68 |
| Average | 2.48 | 40.47 | | 42.95 | 8.70 | 32.80 | | 41.50 | 31.80 | 179.67 | | 211.47 |

**Fraction labelling from the $^{15}$N2-Gln experiment**

**Corrected labelling fraction of metabolite in CNS**

| | Glutamine | | | | Glutamate | | | |
|---|---|---|---|---|---|---|---|---|
| | M+0 | M+1 | M+2 | Mean enrichment | M+0 | M+1 | M+2 | Mean enrichment |
| 1 | 0.01 | 0.00 | 0.99 | 0.99 | 0.10 | 0.90 | | 0.90 |
| 2 | 0.01 | 0.00 | 0.99 | 0.99 | 0.02 | 0.98 | | 0.98 |
| 3 | 0.01 | 0.00 | 0.99 | 0.99 | 0.00 | 1.00 | | 1.00 |
| 4 | 0.01 | 0.00 | 0.99 | 0.99 | 0.06 | 0.94 | | 0.94 |
| 5 | 0.01 | 0.00 | 0.99 | 0.99 | 0.00 | 1.00 | | 1.00 |
| 6 | 0.02 | 0.00 | 0.98 | 0.98 | 0.00 | 1.00 | | 1.00 |
| Average | 0.01 | 0.00 | 0.99 | 0.99 | 0.03 | 0.97 | | 0.97 |

| | GABA | | | | Alanine | | | | Aspartate | | | |
|---|---|---|---|---|---|---|---|---|---|---|---|---|
| | M+0 | M+1 | M+2 | Mean enrichment | M+0 | M+1 | M+2 | Mean enrichment | M+0 | M+1 | M+2 | Mean enrichment |
| 1 | 0.05 | 0.95 | | 0.95 | 0.17 | 0.83 | | 0.83 | 0.09 | 0.91 | | 0.91 |
| 2 | 0.09 | 0.91 | | 0.91 | 0.24 | 0.76 | | 0.76 | 0.08 | 0.92 | | 0.92 |
| 3 | 0.05 | 0.95 | | 0.95 | 0.21 | 0.79 | | 0.79 | 0.18 | 0.82 | | 0.82 |
| 4 | 0.10 | 0.90 | | 0.90 | 0.20 | 0.80 | | 0.80 | 0.13 | 0.87 | | 0.87 |
| 5 | 0.07 | 0.93 | | 0.93 | 0.17 | 0.83 | | 0.83 | 0.24 | 0.76 | | 0.76 |
| 6 | 0.00 | 1.00 | | 1.00 | 0.23 | 0.77 | | 0.77 | 0.17 | 0.83 | | 0.83 |
| Average | 0.06 | 0.94 | | 0.94 | 0.21 | 0.79 | | 0.79 | 0.15 | 0.85 | | 0.85 |

*: fragmented ion used for quantification

### Table 3: Drosophila stocks

| *Drosophila* strains | Genotype | RRID (FlyBase ID) | Stock Number | References |
|---|---|---|---|---|
| w1118 (iso 31) | w[1118] | | gift from J. Roote, University of Cambridge | (Ryder, 2004) PMID: 15238529 |
| repo-Gal4 | w[1118]; P{w[+m*]=GAL4}repo/TM3, Sb[1] | FBti0018692 | Bloomington Stock Center #7415 | (Bailey, 2015) PMID: 26451484 |
| Cyp4g15-Gal4 | w[1118]; P{y[+t7.7] w[+mC]=GMR55B12-GAL4}attP2 | FBti0136853 | Bloomington Stock Center #39103 | (Speder, 2018) PMID: 29299997 |
| moody-Gal4 | w[*]; P{w[+m*]=moody-GAL4.SPG}2 | FBti0212607 | Bloomington Stock Center #90883 | (Vaughen, 2022) PMID: 35961319 |
| daw-Gal4 | | | gift from H. Nakato, University of Minnesota | (Zhu, 2008) PMID: 18171686 |
| nerfin-1-Gal4 | | | gift from A. Kuzin, NIH Bethesda | (Kuzin, 2007) PMID: 17714701 |
| nSyb-Gal4 | y[1] w[*]; P{w[+m*]=nSyb-GAL4.S}3 | FBti0150361 | Bloomington Stock Center #51635 | (Imler, 2019) PMID: 31663851 |
| VGlut1-GAL4 | w[1118]; P{w[+mW.hs]=GawB}VGlut1[OK371] | FBti0076967 | Bloomington Stock Center #26160 | (Mahr, 2006) PMID: 16378756, (Grosjean, 2008) PMID: 18066061 |
| UAS-mCherry RNAi | y[1] sc[*] v[1] sev[21]; P{y[+t7.7] v[+t1.8]=VALIUM20-mCherry.RNAi}attP2 | FBti0143385 | Bloomington Stock Center #35785 | |
| UAS-Gls RNAi | y[1] sc[*] v[1] sev[21]; P{y[+t7.7] v[+t1.8]=TRiP.HMC05223}attP40 | FBti0178998 | Bloomington Stock Center #62216 | (Soares, 2024) PMID: 39047739, (Yadav, 2024) PMID: 39137949 |
| UAS-Gs1 RNAi | y[1] v[1]; P{y[+t7.7] v[+t1.8]=TRiP.HMS02002}attP40 | FBti0149744 | Bloomington Stock Center #40836 | (Pletcher, 2019) PMID: 31036678, (Vernizzi, 2020) PMID: 31941072 |
| UAS-VGlut1 RNAi | y[1] sc[*] v[1] sev[21]; P{y[+t7.7] v[+t1.8]=TRiP.HMS02175}attP40 | FBti0149836 | Bloomington Stock Center #40927 | (Ni, 2019) PMID: 30719975, (Musso, 2021) PMID: 34851668 |
| UAS-VGlut1 RNAi | y[1] sc[*] v[1] sev[21]; P{y[+t7.7] v[+t1.8]=TRiP.HMS02011}attP2 | FBti0149753 | Bloomington Stock Center #40845 | (Ni, 2019) PMID: 30719975, (Musso, 2021) PMID: 34851668 |
| UAS-iGluSnFR | w[1118]; PBac{y[+mDint2] w[+mC]=20XUAS-iGluSnFR.A184A}VK00005/TM3, Sb[1] | FBti0180318 | Bloomington Stock Center #59609 | |
| Gls mutant | y[1] w[*]; TI{GFP[3xP3.cLa]=KozakGAL4}Gls[CR70270-KO-kG4] | FBti0220442 | Bloomington Stock Center #95012 | (Kanca, 2022) PMID: 35723254 |
| Df(Gls) | Df(2R)Exel7121 | FBab0038032 | Bloomington Stock Center #7869 | |
| Gs11 mutant | w[1]; Gs1[1]/CyO | FBal0005224 | Bloomington Stock Center #6248 | (Caggese, 1992) PMID: 1363402 |
| MARCM40A | y,w,hs-Flp,tub-Gal4,UAS-GFPnls / (FM7) ; tub-Gal80, FRT40A / (Cyo) | | gift from M. Amoyel, University College London | (Lee, 1999) PMID: 10197526 |
| FRT40A | y[1] w[*]; P{w[+mC]=tubP-GAL80}LL10 P{ry[+t7.2]=neoFRT}40A/CyO | FBti0002071 | Bloomington Stock Center #5192 | (Xu, 1993) PMID: 8404527 |
| Gls::GFP | y[1] w[67c23]; Mi{PT-GFSTF.0}Gls[MI09647-GFSTF.0]/CyO | FBti0178350 | Bloomington Stock Center #60277 | (Soares, 2024) PMID: 39047739 |
| Eaat1-TG4 | w[1118]; TI{GFP[3xP3.cLa]=CRIMIC.TG4.0}Eaat1[CR01303-TG4.0]/CyO, P{w[+mC]=sChFP}2 | FBti0210329 | Bloomington Stock Center #603635 | (Martelli, 2024) PMID: 38416643 |
| UAS-mCD8::mCherry | w[*]; P{w[+mC]=UAS-mCD8.ChRFP}3 | FBti0115769 | Bloomington Stock Center #27392 | |
| UAS-GFP | w[1118]; P{w[+mC]=UAS-EGFP}5a.2 | FBti0013987 | Bloomington Stock Center #5431 | |
| UAS-RFP | w[1118]; P{w[+mC]=UAS-RFP.W}2 | FBti0129813 | Bloomington Stock Center #30556 | (Wen, 2008) PMID: 18316477 |
| nab-Gal4 | y[*] w[*]; P{w[+mW.hs]=GawB}nab[NP1316] / TM6, P{w[-]=UAS-lacZ.UW23-1}UW23-1 | FBti0034380 | Kyoto DGRC #112622 | (Maurange, 2008) PMID: 18510932 |
| NP6293-Gal4 | y[*] w[*]; P{w[+mW.hs]=GawB}Bsg[NP6293] / CyO, P{w[-]=UAS-lacZ.UW14}UW14 | FBti0036913 | Kyoto DGRC #105188 | (Kanai, 2018) PMID: 29487331 |
| UAS-KK control RNAi | y, w[1118]; P{attP, y[+], w[3']} | | VDRC #60100 | (Green, 2014) PMID: 24577271 |
| UAS-Eaat1 RNAi | y, w[1118]; P{KK100187}VIE-260B | FBst0481090 | VDRC #109401 | (Matsuno, 2019) PMID: 31030182 |

### Table 4: Resources

| Reagent type | Designation | Identifier | Source | Additional Info |
|---|---|---|---|---|
| Antibody | Anti-Deadpan, (rat monoclonal) | ab195173 | Abcam | Dilution 1:500 |
| Antibody | Anti-Miranda, (mouse monoclonal) | | Gift from F. Matsuzaki | Dilution 1:50 |
| Antibody | Anti-Gls, (rabbit polyclonal) | ab93434 | Abcam (discontinued) | Dilution 1:500 |
| Antibody | Anti-Gs1, (rabbit polyclonal) | #11037-2-AP | Proteintech | Dilution 1:500 |
| Antibody | Anti-VGlut1, (mouse monoclonal) | | Gift from B. McCabe | Dilution 1:500 |
| Antibody | Anti-GFP, (rabbit polyclonal) | A-11122 | Invitrogen | Dilution 1:500 |
| Antibody | Anti-Dcp-1, (rabbit polyclonal) | 9578 | Cell Signaling Technology | Dilution 1:500 |
| Antibody | Anti-rat AF488, (goat polyclonal) | #A-11006 | Invitrogen | Dilution 1:1000 |
| Antibody | Anti-rat AF555, (goat polyclonal) | #A-21434 | Invitrogen | Dilution 1:1000 |
| Antibody | Anti-mouse AF488, (goat polyclonal) | #A-11094 | Invitrogen | Dilution 1:1000 |
| Chemical | U-13C-Gln (L-Glutamine-13C5) | 605166 | Sigma-Aldrich | 10 mM |
| Chemical | U-15N-Gln (L-Glutamine-15N2) | 490032 | Sigma-Aldrich | 10 mM |
| Chemical | AF555 Phalloidin | A34055 | Invitrogen | Dilution 1:1000 |
| Chemical | AF647 Phalloidin | A22287 | Invitrogen | Dilution 1:500 |
| Chemical | DAPI | MBD0015 | Sigma-Aldrich | 0.5µg/mL |
| Chemical | Click-iT EdU Alexa Fluor Imaging Kit | C10338 | Invitrogen | |
| Chemical | Vectashield | H-1000-10 | Vector Laboratories | |
| Chemical | Agarose | A9414 | Sigma-Aldrich | |
| Chemical | Triton X-100 | H5141 | Promega | |
| Chemical | NGS | 31873 | Thermo Scientific | |
| Chemical | Penicillin / Streptomycin | 15140122 | Gibco | |
| Chemical | FBS | 12676-029 | Gibco | |
| Chemical | DMSO | D8418 | Sigma-Aldrich | |
| Chemical | PFA 16% | 28906 | Thermo Scientific | |
| Chemical | Glycerol | J64719-AP | Thermo Scientific | |
| Chemical | Nuclease free water | PD092 | Omega Biotek | |
| Chemical | Sucrose (nuclease free) | 419762500 | Thermo Scientific | 250 mM |
| Chemical | 1M Tris (pH 8.0) | 12090-015 | Thermo Scientific | 10 mM |
| Chemical | 1 M KCl | 60142-100ML-F | Sigma-Aldrich | 25 mM |
| Chemical | 1 M MgCl2 | 63069-100ML | Sigma-Aldrich | 5 mM |
| Chemical | 10% Triton-X 100 | 93443-100ML | Sigma-Aldrich | 0.10% |
| Chemical | RNasin Plus | N2615 | Promega | 0.50% |
| Chemical | 50X Protease inhibitor | G6521 | Promega | 1X |
| Chemical | 20 mM DTT | P2325 | Thermo Scientific | 0.1 mM |
| Chemical | 1X PBS | | Crick Media STP | |
| Chemical | 10% BSA (nuclease free) | 126615-25ML | Sigma-Aldrich | 0.50% |
| Chemical | Chloroform | 372978 | Sigma-Aldrich | |
| Chemical | Methanol | 322415 | Sigma-Aldrich | |
| Chemical | Water | 10977-035 | Invitrogen | |
| Chemical | Methanolic m-trifluoromethylphenyl trimethylammonium hydroxide | 79266 | Sigma-Aldrich | |
| Chemical | N,O-Bis(trimethylsilyl)trifluoroacetamide (with 1% trimethylchlorosilane) | B-023 | Supelco | |
| Chemical | Pyridine | 270970 | Sigma-Aldrich | |
| Chemical | Methoxyamine hydrochloride | 89803 | Sigma-Aldrich | |
| Chemical | Pierce$^{TM}$ BCA Protain Assay Kit | 23227 | Fisher Scientific | |
| Chemical | Lauric acid | 167281000 | Fisher Scientific | |
| Chemical | Myristic acid | M3128 | Sigma-Aldrich | |
| Chemical | Palmitic acid | P0500 | Sigma-Aldrich | |
| Chemical | Heptadecanoic acid | H3500 | Sigma-Aldrich | |
| Chemical | Stearic acid | 175366-100G | Sigma-Aldrich | |
| Chemical | Arachidic acid | A3631 | Sigma-Aldrich | |
| Chemical | Heneicosanoic acid | H5149 | Sigma-Aldrich | |
| Chemical | Docosanoic acid | 216941 | Sigma-Aldrich | |
| Chemical | Tetracosanoic acid | L6641 | Sigma-Aldrich | |
| Chemical | (*Z*)-5-tetradecenoic acid | B050241 | BenchChem | |
| Chemical | (*Z*)-7-tetradecenoic acid | BD672807 | BLD Pharm | |
| Chemical | Myristoleic acid | M3525 | Sigma-Aldrich | |
| Chemical | Palmitoleic acid | P9417 | Sigma-Aldrich | |
| Chemical | Oleic acid | O1008 | Sigma-Aldrich | |
| Chemical | Gondoic acid | E3635 | Sigma-Aldrich | |
| Chemical | Erucic acid | E3385 | Sigma-Aldrich | |
| Chemical | Nervonic acid | N1514 | Sigma-Aldrich | |
| Chemical | Linoleic acid | L1376 | Sigma-Aldrich | |
| Chemical | (*Z,Z*)-11,14-eicosadienoic acid | 1192 | Cambridge Bioscience/Matreya | |
| Chemical | (*Z,Z*)-13,16-docosadienoic acid | 20749 | Cayman Chemical | |
| Chemical | Linolenic acid | L2376 | Sigma-Aldrich | |
| Chemical | Palmitic acid-1-$^{13}$C | 292125 | Sigma-Aldrich | |
| Chemical | L-Asparagine | A0884 | Sigma-Aldrich | |
| Chemical | L-Arginine monohydrochloride | A5131 | Sigma-Aldrich | |
| Chemical | L-Alanine | A7627 | Sigma-Aldrich | |
| Chemical | L-Glutamic acid | G1251 | Sigma-Aldrich | |
| Chemical | L-Cystine | C8755 | Sigma-Aldrich | |
| Chemical | L-Cysteine hydrochloride | C1276 | Sigma-Aldrich | |
| Chemical | L-Isoleucine | I2752 | Sigma-Aldrich | |
| Chemical | L-Methionine | M9625 | Sigma-Aldrich | |
| Chemical | L-Tryptophan | T0254 | Sigma-Aldrich | |
| Chemical | L-Tyrosine | T3754 | Sigma-Aldrich | |
| Chemical | L-Threonine | T8625 | Sigma-Aldrich | |
| Chemical | L-Lysine monohydrochloride | L5626 | Sigma-Aldrich | |
| Chemical | L-Leucine | L8000 | Sigma-Aldrich | |
| Chemical | L-Serine | S4500 | Sigma-Aldrich | |
| Chemical | L-Proline | P0380 | Sigma-Aldrich | |
| Chemical | L-Phenylalanine | P2126 | Sigma-Aldrich | |
| Chemical | trans-4-Hydroxy-L-proline | H5534 | Sigma-Aldrich | |
| Chemical | L-Histidine monohydrochloride monohydrate | H8125 | Sigma-Aldrich | |
| Chemical | L-Glutamine | G3126 | Sigma-Aldrich | |
| Chemical | L-Aspartic acid | A9256 | Sigma-Aldrich | |
| Chemical | L-Valine | V0500 | Sigma-Aldrich | |
| Chemical | Glycine | G7126 | Sigma-Aldrich | |
| Chemical | Citric acid monohydrate | C7129 | Sigma-Aldrich | |
| Chemical | ɣ-Aminobutyric acid | A2129 | Sigma-Aldrich | |
| Chemical | β-glycerophosphate disodium salt hydrate | G9422 | Sigma-Aldrich | |
| Chemical | L-Pyroglutamic acid | 83160 | Sigma-Aldrich | |
| Chemical | myo-Inositol | I5125 | Sigma-Aldrich | |
| Chemical | α-D-Glucose | 158968 | Sigma-Aldrich | |
| Chemical | L-(-)-Malic acid | M1000 | Sigma-Aldrich | |
| Chemical | Sodium succinate dibasic hexahydrate | S2378 | Sigma-Aldrich | |
| Chemical | D-Glucose-6-phosphate sodium salt | G7879 | Sigma-Aldrich | |
| Chemical | (+)-Potassium Ds-threo-isocitrate monobasic | 58790 | Sigma-Aldrich | |
| Chemical | scyllo-Inositol | I8132 | Sigma-Aldrich | |
| Chemical | Sodium fumarate dibasic | F1506 | Sigma-Aldrich | |
| Chemical | D-(+)-Trehalose dihydrate | T9531 | Sigma-Aldrich | |
| Chemical | D-Fructose-6-phosphate disodium salt hydrate | F3627 | Sigma-Aldrich | |
| Chemical | Oxaloacetic acid | O7753 | Sigma-Aldrich | |
| Chemical | D-Fructose 1,6-bisphosphate trisodium salt hydrate | F6803 | Sigma-Aldrich | |
| Chemical | α-D-Glucose 1-phoshpate dipotassium salt hydrate | G6750 | Sigma-Aldrich | |
| Chemical | D-(-)-3-Phosphoglyceric acid disodium salt | P8877 | Sigma-Aldrich | |
| Chemical | Phospho(enol)puyruvic acid monopotassium salt | P7127 | Sigma-Aldrich | |
| Chemical | D-2-Phosphoglyceric acid lithium salt | 73885 | Sigma-Aldrich | |
| Chemical | Dihydroxyacetone phosphate dilithium salt | D7137 | Sigma-Aldrich | |
| Chemical | sn-Glycerol 3-phosphate bis(cyclo-hexylammoium) salt | G7886 | Sigma-Aldrich | |
| Chemical | Sodium pyruvate | P2256 | Sigma-Aldrich | |
| Chemical | α-Ketoglutaric acid | K1750 | Sigma-Aldrich | |
| Chemical | L-(+)-Lactic acid | L1750 | Sigma-Aldrich | |
| Chemical | Oxalic acid | 194131-5G | Sigma-Aldrich | |
| Chemical | Urea | U5128-5G | Sigma-Aldrich | |
| Chemical | 2-Picolinic acid | P42800 | Sigma-Aldrich | |
| Chemical | Uracil | A15570.18-50G | Alfa Aesar | |
| Chemical | β-alanine | 146064-25G | Sigma-Aldrich | |
| Chemical | D-erythrose | E7625-250MG | Sigma-Aldrich | |
| Chemical | Creatinine | C4255-10G | Sigma-Aldrich | |
| Chemical | 2-deoxy-D-ribose | 31170-1G-F | Sigma-Aldrich | |
| Chemical | DL-Tartaric acid | T400-25G | Sigma-Aldrich | |
| Chemical | D(+)-xylose | X3877-25G | Sigma-Aldrich | |
| Chemical | L-(+)-arabinose | A91906-1G-A | Sigma-Aldrich | |
| Chemical | D-ribose | R7500-5G | Sigma-Aldrich | |
| Chemical | L-rhamnose monohydrate | R3875-10MG | Sigma-Aldrich | |
| Chemical | L-(-)-fucose | 93183-25MG | Sigma-Aldrich | |
| Chemical | Quinolinic acid | P63204 | Sigma-Aldrich | |
| Chemical | Putrescine dihydrochloride | P5780-5G | Sigma-Aldrich | |
| Chemical | cis-aconitic acid | A3412-1G | Sigma-Aldrich | |
| Chemical | L-ornithine monohydrochloride | O2375-10MG | Sigma-Aldrich | |
| Chemical | 2-methylcitrate | 59464-10MG | Sigma-Aldrich | |
| Chemical | D-fructose | F0127-100G | Sigma-Aldrich | |
| Chemical | D-mannose | CHE1902 | Scientific Laboratory Supplies | |
| Chemical | D-(+)-galactose | G0750-5G | Sigma-Aldrich | |
| Chemical | D-erythrose-4-phosphate sodium salt | E0377-5MG | Sigma-Aldrich | |
| Chemical | Mannitol | M4125-100G | Sigma-Aldrich | |
| Chemical | Sorbitol | S1876-100G | Sigma-Aldrich | |
| Chemical | D-(+)-glucosamine hydrochloride | G4875-25G | Sigma-Aldrich | |
| Chemical | 2-deoxyribose 5-phosphate sodium salt | D3126-25MG | Sigma-Aldrich | |
| Chemical | D-ribose 5-phosphate disodium salt hydrate | R7750-10MG | Sigma-Aldrich | |
| Chemical | D-ribulose 5-phosphate disodium salt | 83899-5MG | Sigma-Aldrich | |
| Chemical | Tryptamine | A11116.09-10G | Alfa Aesar | |
| Chemical | (+)-abscissic acid | 90769-25MG | Sigma-Aldrich | |
| Chemical | D-myo-inositol 3-phosphate sodium salt | 10007778 | Cayman Chemical | |
| Chemical | N-Acetylneuraminic acid | A0812-25MG | Sigma-Aldrich | |
| Chemical | D-sedo-Heptulose 7-phosphate lithium salt | 78832-1MG | Sigma-Aldrich | |
| Chemical | Melatonin | M5250-250MG | Sigma-Aldrich | |
| Chemical | Serotonin hydrochloride | H9523-25MG | Sigma-Aldrich | |
| Chemical | D-lactose monohydrate | L254-25G | Sigma-Aldrich | |
| Chemical | Palatinose hydrate | P2007-10MG | Sigma-Aldrich | |
| Chemical | 2-Methylcitric acid | 6061-96-7 | Cayman Chemical | |
| Chemical | Caffeine | C0750 | Sigma-Aldrich | |
| Chemical | 6-Phosphogluconic acid | P7877 | Sigma-Aldrich | |
| Chemical | D-pantothenic acid hemicalcium salt | P5155 | Sigma-Aldrich | |
| Chemical | Glycerol | G/0600/17 | Fisher Scientific | |
| Chemical | Sucrose | S/8600/53 | Fisher Scientific | |
| Chemical | L-Norleucine | N6877 | Sigma-Aldrich | |
| Chemical | Toluidine Blue O | T3260 | Sigma-Aldrich | |
| Chemical | L-Glutamine-(amine-15N) | 486809-1G | Sigma-Aldrich | |
| Chemical | L-Glutamine-(amide-15N) | 490024-1G | Sigma-Aldrich | |
| Chemical | 2'-Deoxyadenosine monohydrate (13C10, 15N5) | CNLM-3896-CA-5 | Eurisotop | |
| Chemical | Thymidine (methyl-13C) | CLM-3647-PK | Eurisotop | |
| Consumable | Thin wire 0.013 mm (ISAOHM) | 2.4872 | Isabellenhütte | |
| Consumable | Silicone grease | KWIK-SIL | World Precision Instruments, Inc | |
| Consumable | 9-well Pyrex glass plates | 7220-85 | Corning | |
| Consumable | 21-well diagnostic microscope slides | 631-0460 | VWR International | |
| Consumable | Glass coverslips No. 1,5 (24 x 50 mm) | 631-0147 | VWR International | |
| Consumable | Clear nail polish (top coat) | NP90713 | Beauty4Britain | |
| Consumable | RNAzap | R2020-250ML | Sigma-Aldrich | |
| Consumable | 1 mL Dounce | 357538 | Wheaton | |
| Consumable | 20 µm cell strainer | 43-50020-03 | pluriSelect | |
| Consumable | Precellys Metal tubes | P002290-LYSK0-A.0 | Bertin Technologies | |
| Consumable | PTFE O rings (6.07 mm ID x 1.78 mm CS) | BS010PT | Polymax | |
| Consumable | Cell culture dish | 430165 | Corning | |
| Consumable | Positive displacement syringe | 3-000-250 | Drummond | |
| Consumable | Safe-Lock tube | 30120086 | Eppendorf | |
| Consumable | 0.22 µm Filter unit | 8160 | Costar | |
| Consumable | Glass GC vials | 5182-0715 | Agilent Technologies, Inc | |
| Consumable | Glass inserts | 5183-2085 | Agilent Technologies, Inc | |
| Consumable | ITO slides (25 x 25 mm) with a resistivity of 70-100 Ω/sq | 703176 | Sigma-Aldrich | |
| Consumable | J&W DB-5ms GC Column, 30 m, 0.25 mm, 0.25 µm, 7 inch cage | 122-5532 | Agilent Technologies, Inc | |
| Software | FIJI | v2.14.0 | NIH Image | |
| Software | Excel | v16.92 | Microsoft | |
| Software | Illustrator | 2022 | Adobe | |
| Software | Prism 10 | v10.4.1 | GraphPad Software | |
| Software | MassHunter | v10.0 | Agilent Technologies, Inc | |
| Software | IsoCorrectoR | v1.23.0 | | |
| Software | SurfaceLab | v7.3.135519 | IONTOF GmbH | |

### Table 5: Metabolite list with diagnostic ions

**GC-MS peak assignments for 13C5-Gln experiment+A4:F80+A4:F81**

| Assignment | Chemical formula | m/z | Chemical formula for the quantifier ion | Quantifiable labelled C | Note |
|---|---|---|---|---|---|
| Glutamine | **Table 5: Table of metab** | 347 | C13 O3 N2 H31 Si3 | 5 | |
| Glutamate | C5H9NO4 | 246 | C10 O2 N1 H24 Si2 | 4 | fragmented ion |
| GABA | C4H9NO2 | 304 | C12 O2 N1 H30 Si3 | 4 | |
| Alanine | C3H7NO2 | 116 | C5 N1 H14 Si1 | 2 | fragmented ion |
| Pyruvate | C3H4O3 | 174 | C6 O3 N1 H12 Si1 | 3 | |
| Succinate | C4H6O4 | 247 | C9 O4 H19 Si2 | 4 | |
| Fumarate | C4H4O4 | 245 | C9 O4 H17 Si2 | 4 | |
| Aspartate | C4H7NO4 | 232 | C9 O2 N1 H22 Si2 | 3 | fragmented ion |
| Lactate | C3H6O3 | 117 | C5 O1 H13 Si1 | 2 | fragmented ion |

**GC-MS peak assignments for $^{15}N_2$-Gln experiment**

| Assignment | Chemical formula | m/z | Chemical formula for the quantifier ion | Quantifiable labelled N | Note |
|---|---|---|---|---|---|
| Glutamine | C5H10N2O3 | 347 | C13 O3 N2 H31 Si3 | 2 | |
| Glutamate | C5H9NO4 | 348 | C13 O4 N1 H30 Si3 | 1 | |
| GABA | C4H9NO2 | 304 | C12 O2 N1 H30 Si3 | 1 | |
| Alanine | C3H7NO2 | 116 | C5 N1 H14 Si1 | 1 | |
| Aspartate | C4H7NO4 | 334 | C12 O4 N1 H28 Si3 | 1 | |

**OrbiSIMS peak assignments**

| Assignment | Chemical formula | *m/z* | Adduct | Mass Deviation (ppm) |
|---|---|---|---|---|
| Adenine | $C_5H_4N_5^-$ | 134.0473 | $[M-H]^-$ | 0.32 |
| Oleic acid | $C_{18}H_{33}O_2^-$ | 281.2485 | $[M-H]^-$ | -0.24 |
| $^{15}N$ adenine | $C_5H_4N_4{}^{15}N^-$ | 135.0443 | $[M-H]^-$ | 0.08 |
| U-$^{13}C$ adenine | $^{13}C_5H4^{15}N_5^-$ | 144.0491 | $[M-H]^-$ | -0.22 |
| Thymine | $C_5H_5N_2O_2^-$ | 125.0356 | $[M-H]^-$ | -0.23 |
| $^{15}N$ thymine | $C_5H_5{}^{15}NNO_2^-$ | 126.0329 | $[M-H]^-$ | 1.45 |
| Cytosine | $C_4H_4N_3O^-$ | 110.0360 | $[M-H]^-$ | -0.21 |
| $^{15}N$ cytosine | $C_4H_4{}^{15}NN_2O^-$ | 111.0330 | $[M-H]^-$ | -0.45 |
| Guanine | $C_5H_4N_5O^-$ | 150.0421 | $[M-H]^-$ | -0.03 |
| $^{15}N$ guanine | $C_5H_4{}^{15}NN_4O^-$ | 151.0390 | $[M-H]^-$ | -0.83 |
| $^{13}C$ thymine | $C_4{}^{13}CH_5N_2O2^-$ | 126.0390 | $[M-H]^-$ | -0.06 |
| Arginine | $C_6H_{10}ON_4Na^-$ | 177.0754 | $[M+Na-2H]^-$ | -2.01 |
| $^{15}N$ arginine | $C_6H_{10}O^{15}NN_3Na^-$ | 178.0727 | $[M+Na-2H]^-$ | -0.56 |
| Asparagine | $C_4H_5N_2O_2^-$ | 113.0357 | $[M-H]^-$ | 0.13 |
| $^{15}N$ asparagine | $C_4H_5O_2{}^{15}NN^-$ | 114.0326 | $[M-H]^-$ | -1.08 |
| Asparagine | $C_4H_4O_2N_2Na^-$ | 135.0177 | $[M+Na-2H]^-$ | 0.62 |
| $^{15}N$ asparagine | $C_4H_4O_2{}^{15}NNNa^-$ | 136.0147 | $[M+Na-2H]^-$ | 0.18 |
| Aspartic acid | $C_4H_4NO_3^-$ | 114.0196 | $[M-H]^-$ | -0.29 |
| $^{15}N$ aspartic acid | $C_4H_4O_3{}^{15}N^-$ | 115.0167 | $[M-H]^-$ | 0.24 |
| Aspartic acid | $C_4H_3NO_3Na^-$ | 136.0016 | $[M+Na-2H]^-$ | -0.29 |
| $^{15}N$ aspartic acid | $C_4H_3{}^{15}NO_3Na^-$ | 136.9988 | $[M+Na-2H]^-$ | 1.32 |
| Cysteine | $C_3H_4SNO^-$ | 102.0018 | $[M-H]^-$ | -1.03 |
| $^{15}N$ cysteine | $C_3H_4O^{15}NS^-$ | 102.9990 | $[M-H]^-$ | 0.86 |
| Glutamic acid | $C_5H_6NO_3^-$ | 128.0353 | $[M-H]^-$ | 0.12 |
| $^{15}N$ glutamic acid | $C_5H_6O_3{}^{15}N^-$ | 129.0322 | $[M-H]^-$ | -0.83 |
| Glutamic acid | $C_5H_5NO_3Na^-$ | 150.0173 | $[M+Na-2H]^-$ | -0.04 |
| $^{15}N$ glutamic acid | $C_5H_5{}^{15}NNO_3Na^-$ | 165.0174 | $[M+Na-2H]^-$ | -0.13 |
| Glutamine | $C_5H_7N_2O_2^-$ | 127.0513 | $[M-H]^-$ | 0.01 |
| $^{15}N$ glutamine | $C_5H_7O_2{}^{15}NN^-$ | 128.0483 | $[M-H]^-$ | -0.62 |
| Glutamine | $C_5H_6N_2O_2Na^-$ | 149.0333 | $[M+Na-2H]^-$ | 0.29 |
| $^{15}N$ glutamine | $C_5H_6{}^{15}NNO_2Na^-$ | 150.0303 | $[M+Na-2H]^-$ | 0.26 |
| Histidine | $C_6H_6N3O^-$ | 136.0516 | $[M-H]^-$ | -0.25 |
| $^{15}N$ histidine | $C_6H_6O^{15}NN_2^-$ | 137.0486 | $[M-H]^-$ | -0.25 |
| Histidine | $C_6H_5N3ONa^-$ | 158.0336 | $[M+Na-2H]^-$ | -0.13 |
| $^{15}N$ histidine | $C_6H_5{}^{15}NN_2ONa^-$ | 159.0306 | $[M+Na-2H]^-$ | -0.17 |
| Leucine/Isoleucine | $C_6H_{10}NO^-$ | 112.0768 | $[M-H]^-$ | 0.03 |
| $^{15}N$ Leucine/Isoleuc | $C_6H_{10}O^{15}N^-$ | 113.0735 | $[M-H]^-$ | -3.01 |
| Lysine | $C_6H_{11}N_2O^-$ | 127.0877 | $[M-H]^-$ | 0.03 |
| $^{15}N$ lysine | $C_6H_{11}O^{15}NN^-$ | 128.0846 | $[M-H]^-$ | -0.87 |
| Phenylalanine | $C_9H_8NO^-$ | 146.0611 | $[M-H]^-$ | 0.07 |
| $^{15}N$ phenylalanine | $C_9H_8O^{15}N^-$ | 147.0581 | $[M-H]^-$ | -0.16 |
| Phenylalanine | $C_9H_7NONa^-$ | 168.0432 | $[M+Na-2H]^-$ | 0.93 |
| $^{15}N$ phenylalanine | $C_9H_7{}^{15}NONa^-$ | 169.0402 | $[M+Na-2H]^-$ | 0.63 |
| Serine | $C_3H_3O_2NNa^-$ | 108.0066 | $[M+Na-2H]^-$ | -0.72 |
| $^{15}N$ serine | $C_3H_3O_2{}^{15}NNa^-$ | 109.0037 | $[M+Na-2H]^-$ | 0.15 |
| Threonine | $C_4H_6NO_2^-$ | 100.0403 | $[M-H]^-$ | -1.21 |
| $^{15}N$ threonine | $C_4H_6O_2{}^{15}N^-$ | 101.0376 | $[M-H]^-$ | 1.17 |
| Tryptophan | $C_{11}H_9N_2O^-$ | 185.0721 | $[M-H]^-$ | 0.13 |
| $^{15}N$ tryptophan | $C_{11}H_9O^{15}NN^-$ | 186.0691 | $[M-H]^-$ | 0.28 |
| Tryptophan | $C_{11}H_8N_2ONa^-$ | 207.0541 | $[M+Na-2H]^-$ | 0.34 |
| $^{15}N$ tryptophan | $C_{11}H_8{}^{15}NNONa^-$ | 208.0502 | $[M+Na-2H]^-$ | -3.73 |
| Tyrosine | $C_9H_8NO_2^-$ | 162.0561 | $[M-H]^-$ | 0.22 |
| $^{15}N$ tyrosine | $C_9H_8O_2{}^{15}N^-$ | 163.0531 | $[M-H]^-$ | -0.15 |
| Tyrosine | $C_9H_7NO_2Na^-$ | 184.0380 | $[M+Na-2H]^-$ | -0.13 |
| $^{15}N$ tyrosine | $C_9H_7{}^{15}NO_2Na^-$ | 185.0348 | $[M+Na-2H]^-$ | -0.99 |
